# Virological characteristics of the SARS-CoV-2 Omicron BA.2.75

**DOI:** 10.1101/2022.08.07.503115

**Authors:** Akatsuki Saito, Tomokazu Tamura, Jiri Zahradnik, Sayaka Deguchi, Koshiro Tabata, Izumi Kimura, Jumpei Ito, Hesham Nasser, Mako Toyoda, Kayoko Nagata, Keiya Uriu, Yusuke Kosugi, Shigeru Fujita, Daichi Yamasoba, Maya Shofa, MST Monira Begum, Yoshitaka Oda, Rigel Suzuki, Hayato Ito, Naganori Nao, Lei Wang, Masumi Tsuda, Kumiko Yoshimatsu, Yuki Yamamoto, Tetsuharu Nagamoto, Hiroyuki Asakura, Mami Nagashima, Kenji Sadamasu, Kazuhisa Yoshimura, Takamasa Ueno, Gideon Schreiber, Akifumi Takaori-Kondo, The Genotype to Phenotype Japan (G2P-Japan) Consortium, Kotaro Shirakawa, Hirofumi Sawa, Takashi Irie, Kazuo Takayama, Keita Matsuno, Shinya Tanaka, Terumasa Ikeda, Takasuke Fukuhara, Kei Sato

**Affiliations:** Department of Veterinary Science, Faculty of Agriculture, University of Miyazaki, Miyazaki, Japan; Graduate School of Medicine and Veterinary Medicine, University of Miyazaki, Miyazaki, Japan; Center for Animal Disease Control, University of Miyazaki, Miyazaki, Japan; Department of Microbiology and Immunology, Faculty of Medicine, Hokkaido University, Sapporo, Japan.; Department of Biomolecular Sciences, Weizmann Institute of Science, Rehovot, Israel; Center for iPS Cell Research and Application (CiRA), Kyoto University, Kyoto, Japan; Division of Molecular Pathobiology, International Institute for Zoonosis Control, Hokkaido University, Sapporo, Japan; Division of Systems Virology, Department of Microbiology and Immunology, The Institute of Medical Science, The University of Tokyo, Tokyo, Japan; Division of Molecular Virology and Genetics, Joint Research Center for Human Retrovirus infection, Kumamoto University, Kumamoto, Japan; Department of Clinical Pathology, Faculty of Medicine, Suez Canal University, Ismailia, Egypt; Division of Infection and immunity, Joint Research Center for Human Retrovirus infection, Kumamoto University, Japan; Department of Hematology and Oncology, Graduate School of Medicine, Kyoto University, Kyoto, Japan; Graduate School of Medicine, The University of Tokyo, Tokyo, Japan; Faculty of Medicine, Kobe University, Kobe, Japan; Department of Cancer Pathology, Faculty of Medicine, Hokkaido University, Sapporo, Japan; Division of International Research Promotion, International Institute for Zoonosis Control, Hokkaido University, Sapporo, Japan; Institute for Chemical Reaction Design and Discovery (WPI-ICReDD), Hokkaido University, Sapporo, Japan; Institute for Genetic Medicine, Hokkaido University, Sapporo, Japan; HiLung Inc., Kyoto, Japan; Tokyo Metropolitan Institute of Public Health, Tokyo, Japan; One Health Research Center, Hokkaido University, Sapporo, Japan.; Institute of Biomedical and Health Sciences, Hiroshima University, Hiroshima, Japan.; AMED-CREST, Japan Agency for Medical Research and Development (AMED), Tokyo, Japan; International Collaboration Unit, International Institute for Zoonosis Control, Hokkaido University, Sapporo, Japan; Division of Risk Analysis and Management, International Institute for Zoonosis Control, Hokkaido University, Sapporo, Japan; Laboratory of Virus Control, Research Institute for Microbial Diseases, Osaka University, Suita, Japan.; International Research Center for Infectious Diseases, The Institute of Medical Science, The University of Tokyo, Tokyo, Japan; International Vaccine Design Center, The Institute of Medical Science, The University of Tokyo, Tokyo, Japan; Graduate School of Frontier Sciences, The University of Tokyo, Kashiwa, Japan; Collaboration Unit for Infection, Joint Research Center for Human Retrovirus infection, Kumamoto University, Kumamoto, Japan; CREST, Japan Science and Technology Agency, Kawaguchi, Japan

**Keywords:** SARS-CoV-2, COVID-19, Omicron, BA.2.75, transmissibility, immune resistance, antiviral drug resistance, pathogenicity

## Abstract

SARS-CoV-2 Omicron BA.2.75 emerged in May 2022. BA.2.75 is a BA.2 descendant but is phylogenetically different from BA.5, the currently predominant BA.2 descendant. Here, we showed that the effective reproduction number of BA.2.75 is greater than that of BA.5. While the sensitivity of BA.2.75 to vaccination- and BA.1/2 breakthrough infection-induced humoral immunity was comparable to that of BA.2, the immunogenicity of BA.2.75 was different from that of BA.2 and BA.5. Three clinically-available antiviral drugs were effective against BA.2.75. BA.2.75 spike exhibited a profound higher affinity to human ACE2 than BA.2 and BA.5 spikes. The fusogenicity, growth efficiency in human alveolar epithelial cells, and intrinsic pathogenicity in hamsters of BA.2.75 were comparable to those of BA.5 but were greater than those of BA.2. Our multiscale investigations suggest that BA.2.75 acquired virological properties independently of BA.5, and the potential risk of BA.2.75 to global health is greater than that of BA.5.

## Introduction

By the end of 2021, five SARS-CoV-2 variants-of-concern (VOCs) were classified by the WHO (WHO, 2022). These are the Alpha [also known as lineage B.1.1.7 based on the PANGO classification (https://cov-lineages.org); clade 20I based on the Nextstrain classification (https://nextstrain.org)], Beta (lineage B.1.351; clade 20H), Gamma (lineage P.1; clade 20J), Delta (lineages B.1.617.2 and AY; clades 21I and 21J), and Omicron (lineages B.1.1.529 and BA; clade 21K) variants. Since these five VOCs are phylogenetically unrelated to each other, SARS-CoV-2 evolution until the end of 2021 was posed by the antigenic shift. At the beginning of 2022, Omicron BA.1 variant (clade 21K) outcompeted the other variants and spread globally. Thereafter, BA.2 (clade 21L) and BA.4/5 (clades 22A and 22B) continuously emerged from South Africa, while BA.2.12.1 (clade 22C) emerged in the USA. As of the beginning of August 2022, Omicron BA.5 (clade 22B) is the most predominant SARS-CoV-2 variant in the world. In contrast to the five VOCs detected in 2021, the Omicron subvariants are phylogenetically related. Therefore, the evolution of SARS-CoV-2 Omicron subvariants since the end of 2021 is posed by the antigenic drift.

Newly emerging SARS-CoV-2 variants need to be carefully and rapidly assessed for a potential increase in their growth efficiency in the human population [i.e., relative effective reproduction number (R_e_)], their evasion from antiviral immunity, and their pathogenicity. Resistance to antiviral humoral immunity can be mainly determined by substitutions in the spike (S) protein. For instance, Omicron BA.1 (Cao et al., 2021; Cele et al., 2021; Dejnirattisai et al., 2022; Garcia-Beltran et al., 2021; Liu et al., 2021; Meng et al., 2022; Planas et al., 2021; Takashita et al., 2022a; VanBlargan et al., 2022) , BA.2 (Bruel et al., 2022; Takashita et al., 2022b; Yamasoba et al., 2022c), and BA.5 (Arora et al., 2022; Cao et al., 2022; Gruell et al., 2022; Hachmann et al., 2022; Khan et al., 2022; Kimura et al., 2022c; Lyke et al., 2022; Qu et al., 2022; Tuekprakhon et al., 2022; Wang et al., 2022; Yamasoba *et al*., 2022c) exhibit profound resistance to neutralizing antibodies induced by vaccination, natural SARS-CoV-2 infection, and therapeutic monoclonal antibodies. Particularly, newly spreading SARS-CoV-2 variants tend to be resistant to the humoral immunity induced by the infection with prior variant; for instance, BA.2 is resistant to BA.1 breakthrough infection sera (Qu *et al*., 2022; Tuekprakhon *et al*., 2022; Yamasoba et al., 2022b), and BA.5 is resistant to BA.2 breakthrough infection sera (Hachmann *et al*., 2022; Kimura *et al*., 2022c; Wang *et al*., 2022). Therefore, acquiring immune resistance to previously dominant variant is a key factor in outcompeting previous variants, thereby obtaining relatively increased R_e_ compared to the previously dominant variant. Viral pathogenicity is also closely associated with the phenotype of viral S protein. Particularly, we have proposed that the fusogenicity of viral S protein in *in vitro* cell cultures is associated with viral pathogenicity *in vivo* (Kimura *et al*., 2022c; Saito et al., 2022; Suzuki et al., 2022; Yamasoba *et al*., 2022b).

As mentioned above, major SARS-CoV-2 phenotypes can be defined by the function of the viral S protein. SARS-CoV-2 S protein bears two major domains, receptor binding domain (RBD) and N-terminal domain (NTD) [reviewed in (Harvey et al., 2021; Mittal et al., 2022)]. RBD is crucial for the binding to human angiotensin-converting enzyme 2 (ACE2) receptor for the cell attachment and entry, and therefore, this domain has been considered a major target for neutralizing antibodies to block viral infection [reviewed in (Barnes et al., 2020; Harvey *et al*., 2021; Jackson et al., 2022)]. On the other hand, NTD is an immunodominant domain that can be recognized by antibodies, and some antibodies targeting NTD potentially neutralize viral infection (Cerutti et al., 2021; Chi et al., 2020; Liu et al., 2020; Lok, 2021; McCallum et al., 2021; Suryadevara et al., 2021; Voss et al., 2021), despite our limited understanding of its virological function.

The Omicron BA.2.75 variant, a new BA.2 subvariant, was first detected in India in May 2022 (WHO, 2022). Because an early preliminary investigation suggested the potential increase in the relative R_e_ value of BA.2.75 compared to BA.5 and the original BA.2 (GitHub, 2022), BA.2.75 has been flagged as the most concerning variant that can potentially outcompete BA.5 and be the next predominant variant in the future. In fact, on July 19, 2022, the WHO classified this variant as a VOC lineage under monitoring (VOC-LUM) together with the other BA.2 subvariants, including BA.5, which bear the substitution at the L452 residue in their S proteins (WHO, 2022). On July 23, 2022, Nextstrain (https://nextstrain.org) classified BA.2.75 as a new clade, 22D. Compared to the BA.2 S, BA.4/5 bears four mutations in its S protein (Kimura *et al*., 2022c; Yamasoba *et al*., 2022b). On the other hand, the majority of BA.2.75 S bears nine mutations: K147E, W152R, F157L, I210V, and G257S substitutions are located in the NTD, while D339H, G446S, N460K, and R493Q substitutions are located in the RBD. The mutation number in the BA.2.75 S is larger than that in the BA.4/5 S, and notably, some of the substitutions detected in the BA.2.75 S show the signs of convergent evolution (Zahradnik et al., 2022). These notions raise the possibility that the phenotype of BA.2.75 S is critically different from previous BA.2 subvariants. In fact, we have recently revealed that the S protein of BA.2.75 exhibits different sensitivity towards several therapeutic monoclonal antibodies from those of BA.2 and BA.5 (Yamasoba et al., 2022a). However, the virological phenotype of BA.2.75, including its R_e_, potential evasion from antiviral humoral immunity, sensitivity to currently recommended antiviral small compounds, virological properties of its S protein, and intrinsic pathogenicity remains unclear. Here, we elucidate the features of newly emerging SARS-CoV-2 Omicron BA.2.75 subvariant.

## Results

### Epidemics of BA.2.75 in India

As of the beginning of August 2022, the Omicron BA.5 variant is predominant in the world and is outcompeting the BA.2 variant. However, a novel BA.2 subvariant, BA.2.75, emerged and rapidly spread in India since May 2022. Although BA.2.75 and BA.5 (and BA.4) belong to the BA.2 subvariant clade, BA.2.75 is phylogenetically distinct from the BA.4/5 clade (**Figure 1A**). Compared to BA.2, BA.2.75 harbors 14 amino acid substitutions, including nine substitutions in the S protein (**Figures 1B** **and S1A**). Of these, only one revertant mutation (S:R493Q) is shared with BA.5. In India, BA.5 and BA.2.75 spread in different regions each other: BA.5 spreads in the south part including Tamil Nadu and Telangana states, while BA.2.75 spreads the other parts including Himachal Pradesh, Odisha, Haryana, Rajasthan, and Maharashtra states (**Figures 1C and 1D**). To compare the relative R_e_ between BA.5 and BA.2.75 in India with adjusting the regional differences, we constructed a Bayesian hierarchical model that can estimate both state-specific R_e_ values and the value averaged in India (**Figures 1E** **and S1B and Table S1**). The R_e_ value of BA.5 is 1.19-fold higher than that of BA.2 [95% credible interval (CI): 1.14–1.24] on average in India (**Figure 1E**). This value is comparative to the relative R_e_ value of BA.5 in South Africa (1.21) estimated in our recent study (Kimura *et al*., 2022c). Of note, the R_e_ value of BA.2.75 is 1.34-fold higher than that of BA.2 (95% CI: 1.29–1.38), and the R_e_ value of BA.2.75 is 1.13-fold higher than that of BA.5 (95% CI: 1.06–1.20) (**Figures 1E** **and S1C**). Furthermore, in the Indian states analyzed, where both BA.5 and BA.2.75 are dominant, such as Telangana and Tamil Nadu (for BA. 5-dominant states) and Odisha, Haryana, Rajasthan, and Maharashtra (for BA.2.75-dominant states), the R_e_ value of BA.2.75 was greater than that of BA.5 (**Figures S1B and S1C**). Together, our data suggest that BA.2.75 bears the potential to spread more rapidly than BA.5 and will be predominant in some regions including India in the near future.

**Figure 1.**
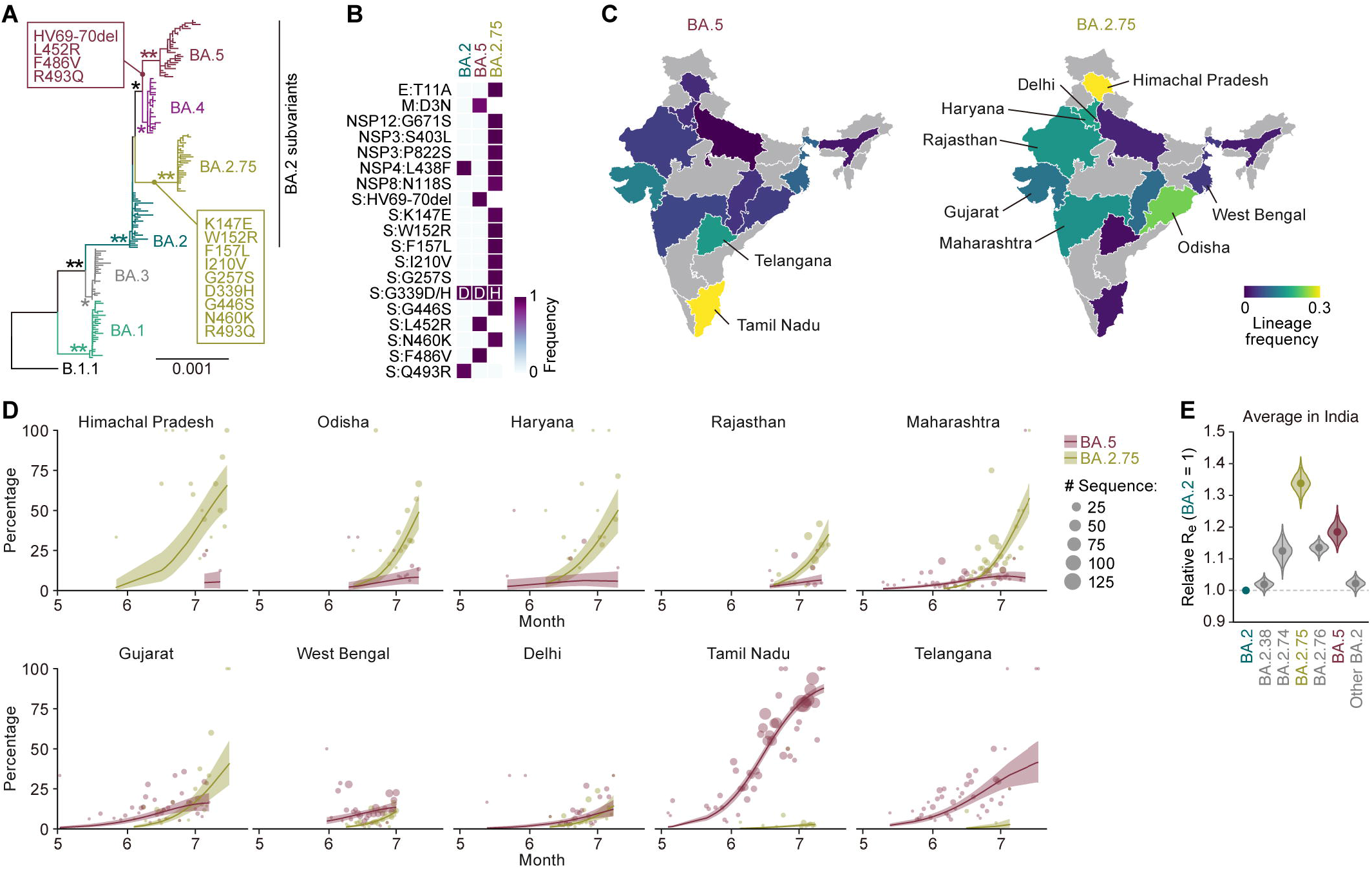
Epidemics of BA.2.75 in India. (**A**) A maximum likelihood tree of Omicron sublineages. Sequences of BA.1–BA.5 sampled from South Africa and BA.2.75 are included. The mutations acquired in the S protein of BA.2.75 are indicated in the panel. Note that R493Q is a reversion [i.e., back mutation from the BA.1–BA.3 lineages (R493) to the B.1.1 lineage (Q493)]. Bootstrap values, *, ≥ 0.8; **, ≥ 0.95. (**B**) Amino acid differences among BA.2, BA.2.75, and BA.5. Heatmap color indicates the frequency of amino acid substitutions. (**C**) Lineage frequencies of BA.5 (left) and BA.2.75 (right) in each Indian state. SARS-CoV-2 sequences collected from June 15, 2022, to July 15, 2022, were analyzed. (**D**) Epidemic dynamics of SARS-CoV-2 lineages in Indian states. Results for BA.2.75 and BA.5 are shown. The observed daily sequence frequency (dot) and the dynamics (posterior mean, line; 95% CI, ribbon) are shown. The dot size is proportional to the number of sequences. (**E**) Estimated relative R_e_ of each viral lineage, assuming a fixed generation time of 2.1 days. The R_e_ value of BA.2 is set at 1. The posterior (violin), posterior mean (dot), and 95% CI (line) are shown. The average values across India estimated by a Bayesian hierarchical model are shown, and the state-specific R_e_ values are shown in Figure S1B. The dynamics of the top seven predominant lineages in India were estimated. BA.5 sublineages are summarized as “BA.5”, and non-predominant BA.2 sublineages are summarized as “other BA.2”. See also Figure S1 and Table S1.

### Sensitivity of BA.2.75 to antiviral humoral immunity and antiviral drugs

Recent studies, including ours, showed that newly emerging Omicron subvariants such as BA.5 exhibit higher resistance to the humoral immunity induced by vaccination and natural infections with prior SARS-CoV-2 variants including BA.1 and BA.2 (Hachmann *et al*., 2022; Kimura *et al*., 2022c; Wang *et al*., 2022). Additionally, we have recently demonstrated that BA.2.75 is more resistant to a therapeutic monoclonal antibody, bebtelovimab, compared to BA.2 and BA.5 (Yamasoba *et al*., 2022a). To investigate the sensitivity of BA.2.75 to antiviral humoral immunity, we prepared pseudoviruses bearing the S proteins of D614G-bearing ancestral B.1.1, BA.2, BA.5 and BA.2.75. Human sera were collected from vaccinated and infected individuals (listed in **Table S2**). The 2-dose vaccine sera were ineffective against all Omicron subvariants tested, including BA.2.75 (**Figure 2A**). Although BA.5 was significantly more resistant to 3-dose vaccine sera than BA.2, which is consistent with previous studies (Hachmann *et al*., 2022; Kimura *et al*., 2022c; Wang *et al*., 2022), the sensitivity of BA.2.75 to these sera was comparable to that of BA.2 (**Figures 2B and 2C**). We then assessed the sensitivity of BA.2.75 to the convalescent sera from individuals who were infected with BA.1 and BA.2 after 2-dose or 3-dose vaccination (i.e., breakthrough infection). Similar to the previous reports including ours (Hachmann *et al*., 2022; Kimura *et al*., 2022c; Wang *et al*., 2022), BA.5 exhibited significant resistance to breakthrough infection sera compared to BA.2, while the sensitivity of BA.2.75 to these sera was comparable to that of BA.2 (**Figures 2D and 2E**). These results suggest that BA.2.75 is not resistant to the humoral immunity induced by vaccination and the infection with prior Omicron subvariants including BA.1 and BA.2. Since the Delta variant emerged and caused a huge surge of infection in India in the middle of 2021 (Mlcochova et al., 2021), it is hypothesized that BA.2.75 evades the immunity induced by Delta. To address this possibility, we used Delta infection sera. However, the sensitivity of all Omicron subvariants tested, including BA.2.75, to Delta infection sera was similar (**Figure 2F**), implying that previous Delta infection is not associated with the emergence of BA.2.75 in India.

**Figure 2.**
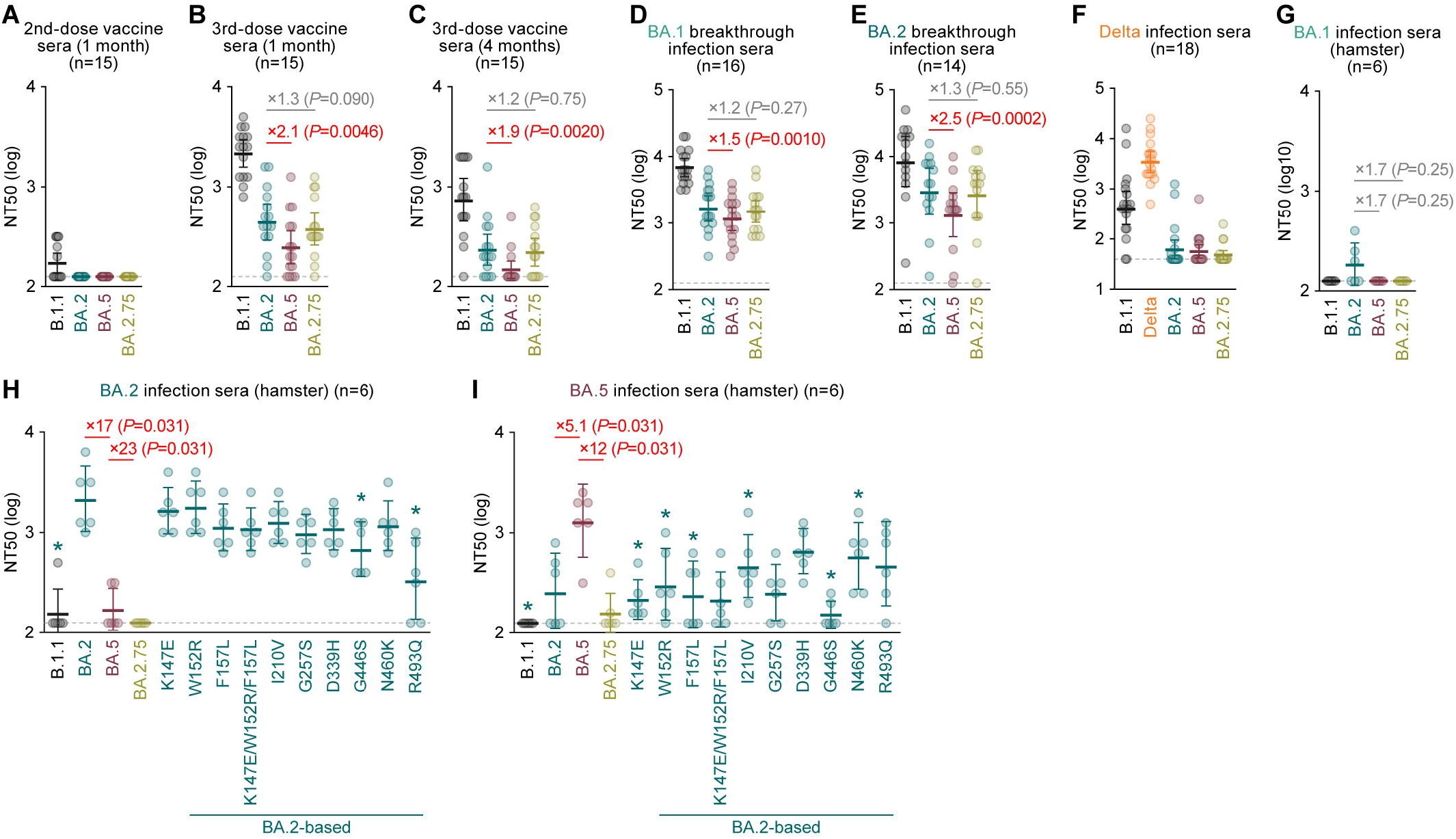
Immune resistance of BA.2.75. Neutralization assays were performed with pseudoviruses harboring the S proteins of B.1.1 (the D614G-bearing ancestral virus), BA.1, BA.2, BA.2.75. Delta pseudovirus is included only in the experiment shown in F. The BA.2 S-based derivatives are included in H and I. The following sera were used. (**A–C**) BNT162b2 vaccine sera (15 donors) collected at 1 month after 2nd-dose vaccination (A), 1 month after 3rd-dose vaccination (B), and 4 months after 3rd-dose vaccination (**C**). (**D**) Convalescent sera from fully vaccinated individuals who had been infected with BA.1 after full vaccination (16 2-dose vaccinated donors). (**E**) Convalescent sera from fully vaccinated individuals who had been infected with BA.2 after full vaccination (9 2-dose vaccinated and 5 3-dose vaccinated. 14 donors in total). (**F**) Convalescent sera from unvaccinated individuals who had been infected with Delta (18 donors). (**G–I**) Sera from hamsters infected with BA.1 (6 hamsters) (**G**), BA.2 (6 hamsters) (**H**), and BA.5 (6 hamsters) (**I**). Assays for each serum sample were performed in triplicate to determine the 50% neutralization titer (NT50). Each dot represents one NT50 value, and the geometric mean and 95% CI are shown. The numbers in the panels indicate the fold change resistance versus BA.2 (**B–E, G** and **H**) or BA.5 (**I**). The horizontal dashed line indicates the detection limit (120-fold in other than **F**, 40-fold in **F**). Statistically significant differences were determined by two-sided Wilcoxon signed-rank tests. The *P* values versus BA.2 (**B–E, G and H**) or BA.5 (**I**) are indicated in the panels. For the BA.2 derivatives and B.1.1 (**H and I**), statistically significant differences versus BA.2 (*P* < 0.05) are indicated with asterisks. Information on the vaccinated/convalescent donors is summarized in **Table S2**. See also **Table S2**.

To further address the difference in immunogenicity among Omicron subvariants, we used the sera obtained from infected hamsters at 16 days postinfection (d.p.i., i.e., after recovery) (Kimura *et al*., 2022c; Suzuki *et al*., 2022; Yamasoba *et al*., 2022b). While BA.1 infection hamster sera were ineffective against BA.2, BA.5 and BA.2.75 (**Figure 2G**), both BA.5 (17-fold, *P*=0.031 by Wilcoxon signed-rank test) and BA.2.75 (23-fold, *P*=0.031 by Wilcoxon signed-rank test) exhibited significant resistance to BA.2 infection hamster sera than BA.2 (**Figure 2H**). These results suggest that the immunogenicity of BA.5 and BA.2.75 is different from BA.2. Notably, BA.2 (5.1-fold, *P*=0.031 by Wilcoxon signed-rank test) and BA.2.75 (12-fold, *P*=0.031 by Wilcoxon signed-rank test) exhibited significant resistance to BA.5 infection hamster sera (**Figure 2I**). These results suggest that the immunogenicity of BA.5 and BA.2.75 is also different. To identify the substitutions responsible for the different immunogenicity of BA.2.75 S from BA.2 S and BA.5 S, we prepared the BA.2 S-based derivatives that bear respective BA.2.75 substitutions. The neutralization assay using BA.2-infected hamster sera showed that the G446S and R493Q substitutions contribute to the resistance of BA.2.75 to BA.2-induced immunity (**Figure 2H**). Because the R493Q substitution is shared with BA.5 (**Figures 1B** **and S1A**), it can be suggested that this substitution contributes to the resistance of BA.5 to BA.2-induced immunity (**Figure 2H**). In the case of BA.5-infected hamster sera, multiple substitutions, including the K147E, W152R, F157L, I210V, G446S and N460K, associated with the resistance of BA.2.75 to BA.5-induced immunity (**Figure 2I**).

To evaluate the sensitivity of BA.2.75 to three antiviral drugs, Remdesivir, EIDD-1931 (an active metabolite of Molnupiravir) and Nirmatrelvir (also known as PF-07321332), we used a clinical isolate of BA.2.75 (strain TY41-716; GISAID ID: EPI_ISL_13969765). As controls, we also used clinical isolates of B.1.1 (strain TKYE610670; GISAID ID: EPI_ISL_479681) (Suzuki *et al*., 2022), BA.2 (strain TY40-385; GISAID ID: EPI_ISL_9595859) (Kimura *et al*., 2022c), BA.5 (strain TKYS14631; GISAID ID: EPI_ISL_12812500) (Tamura et al., 2022). These viruses were inoculated into human airway organoids (AO), a physiologically relevant model (Sano et al., 2022), and treated with three antiviral drugs. As shown in **Table 1** and **Figure S2A**, Remdesivir had a stronger antiviral effect (EC_50_=0.63 μM) against B.2.75 than other variants, B.1.1, BA.2 and BA.5. EIDD-1931 inhibited BA.2 and BA.2.75 (EC_50_=0.02 μM and 0.08 μM, respectively) more potently than B.1.1 and BA.5 (EC_50_=0.24 μM and 0.21 μM, respectively). For Nirmatrelvir, no differences in antiviral efficacy were observed between four variants (EC_50_=0.84 μM, 0.85 μM, 0.63 μM and 0.81 μM for B.1.1, BA.2, BA.5 and BA.2.75, respectively). Altogether, it is suggested that all three drugs exhibit antiviral effects against BA.2.75, and particularly, EIDD-1931 is effective against BA.2.75.

**Table 1.** Effects of three antiviral drugs against BA.2.75 in AO

|  | EC<sub>50</sub> (μM) |  |  |  | EC<sub>50</sub> ratio |  |  |  |  |
| --- | --- | --- | --- | --- | --- | --- | --- | --- | --- |
|  | B.1.1 | BA.2 | BA.5 | BA.2.75 | BA.5/BA.2 | BA.2.75/BA.2 | BA.2/B.1.1 | BA.5/B.1.1 | BA.2.75/B.1.1 |
| Remdesivir | 1.08 | 1.89 | 1.31 | 0.63 | 0.70 | 0.34 | 1.75 | 1.21 | 0.59 |
| EIDD-1931 | 0.24 | 0.02 | 0.21 | 0.08 | 8.82 | 3.53 | 0.10 | 0.89 | 0.36 |
| Nirmatrelvir | 0.84 | 0.85 | 0.63 | 0.81 | 0.74 | 0.95 | 1.02 | 0.75 | 0.97 |

### Virological characteristics of BA.2.75 S *in vitro*

To investigate the virological properties of BA.2.75 S, we measured the pseudovirus infectivity. As shown in **Figure 3A**, the pseudovirus infectivity of BA.2.75 was significantly (12.5-fold) higher than that of BA.2. To assess the association of TMPRSS2 usage with the increased pseudovirus infectivity of BA.2.75, we used both HEK293-ACE2/TMPRSS2 cells and HEK293-ACE2 cells, on which endogenous surface TMPRSS2 is undetectable (Yamasoba *et al*., 2022b), as target cells. Consistent with our recent study (Kimura *et al*., 2022c), the fold increase in pseudovirus infectivity of BA.5 caused by TMPRSS2 expression on the target cells was not observed (**Figure S3A**). Similarly, the infectivity of BA.2.75 pseudovirus was not increased by TMPRSS2 expression (**Figure S3A**), suggesting that TMPRSS2 is not associated with an increase in pseudovirus infectivity of BA.2.75. To determine the substitutions that are responsible for the increased pseudovirus infectivity of BA.2.75, we used a series of BA.2 derivatives that bears the BA.2.75-specific substitutions. Three substitutions in the NTD, K147E, F157L, and I210V, and two substitutions in the RBD, N460K and R493Q, significantly increased infectivity (**Figure 3A**). Notably, the N460K substitution increased infectivity by 44-fold (**Figure 3A**). On the other hand, a substitution in the NTD, W152R, significantly (8.9-fold) decreased infectivity (**Figure 3A**). The BA.2 derivative bearing the three substitutions in the NTD in close proximity to each other, K147E, W152R and F157L, exhibited comparable infectivity to BA.2 (**Figure 3A**).

**Figure 3.**
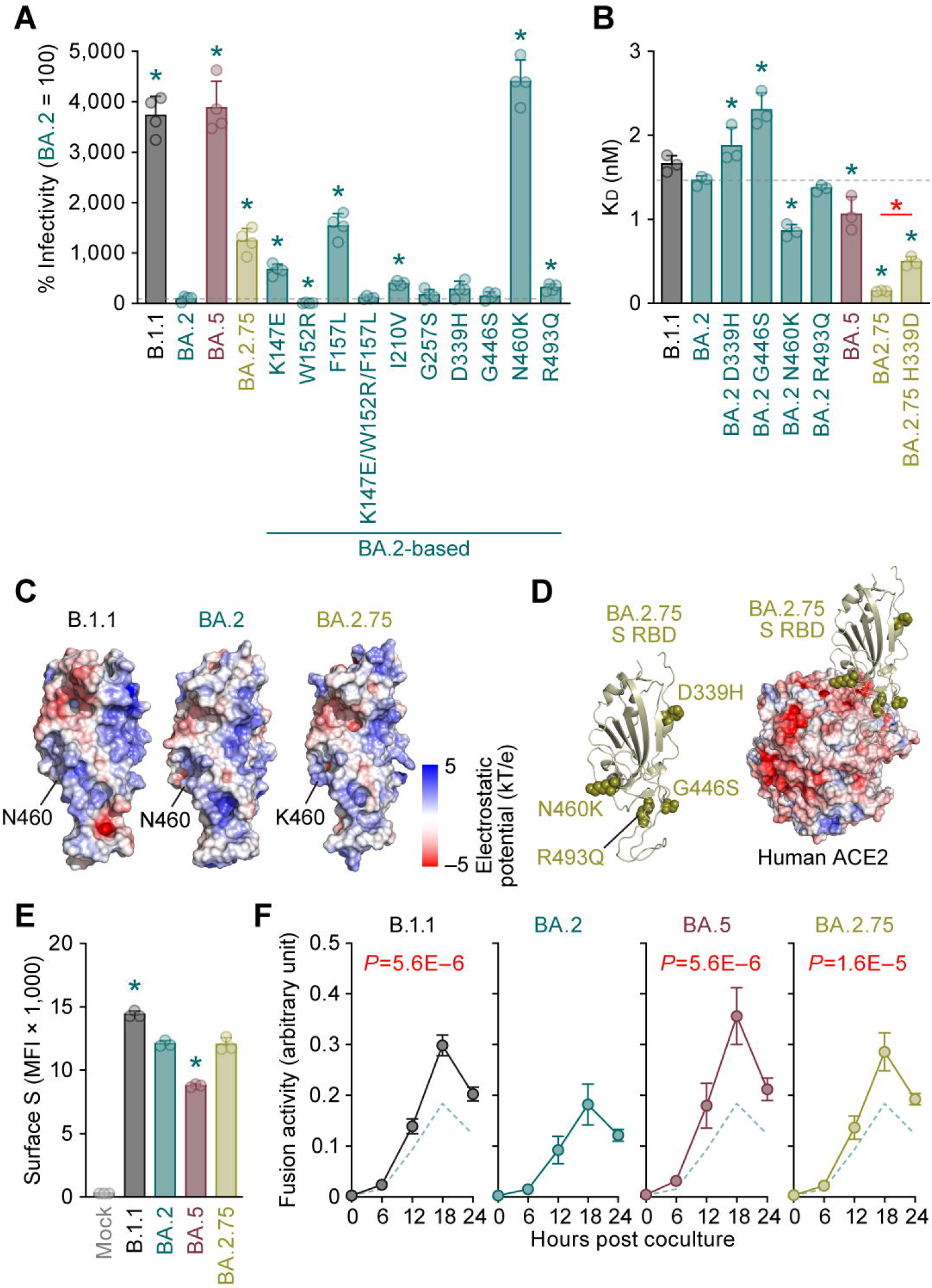
Virological features of BA.2.75 S *in vitro*. (**A**) Pseudovirus assay. The percent infectivity compared to that of the virus pseudotyped with the BA.2 S protein are shown. (**B**) Binding affinity of the RBD of SARS-CoV-2 S protein to ACE2 by yeast surface display. The K_D_ value indicating the binding affinity of the RBD of the SARS-CoV-2 S protein to soluble ACE2 when expressed on yeast is shown. (**C**) Electrostatic potential of B.1.1 S RBD (PDB: 6M17) (Yan *et al*., 2020), BA.2 S RBD (PDB: 7UB0) (Stalls *et al*., 2022) and BA.2.75 S RBD. The structure of BA.2.75 S RBD was prepared using AlphaFold2 (Mirdita *et al*., 2022). Electrostatic potential surface depictions calculated by PDB2PQR tool (Dolinsky *et al*., 2007) with the positions of BA.2.75 characteristic mutations. The scale bar shows the electrostatic charge [kT/e]. (**D**) The binding of BA.2.75 S RBD and human ACE2 (PDB: 6M17) (Yan et al., 2020). Left, the four substitutions in BA.2.75 S RBD compared to BA.2 S RBD are highlighted. Right, binding of BA.2.75 S RBD (top) and human ACE2 (bottom). The electrostatic potential surface of human ACE2 is shown. (**E and F**) S-based fusion assay. (**E**) S protein expression on the cell surface. The summarized data are shown. (F) S-based fusion assay in Calu-3 cells. The recorded fusion activity (arbitrary units) is shown. The dashed green line indicates the results of BA.2. Assays were performed in quadruplicate (A and F) or triplicate (B and E), and the presented data are expressed as the average ± SD. In A and B, the dashed horizontal lines indicated the value of BA.2. In A, B and E, each dot indicates the result of an individual replicate. In A, B and E, statistically significant differences between BA.2 and other variants (*, *P* < 0.05) were determined by two-sided Student’s *t* tests. In F, statistically significant differences between BA.2 and other variants across timepoints were determined by multiple regression. The FWERs calculated using the Holm method are indicated in the figures. See also Figure S3.

To decipher the binding properties of BA.2.75 S RBD to human ACE2 and the role of each substitution, we measured the ACE2 binding affinity of the S RBDs of BA.2.75 as well as those of BA.2 derivatives bearing D339H, G446S, N460K and R493Q substitutions by an enhanced surface display system (Zahradnik et al., 2021a). Intriguingly, the BA.2.75 S RBD showed a strongly tight binding with 146 ± 6 pM affinity (**Figure 3B**). Out of the four BA.2-based derivatives, only the BA.2 N460K substitution exhibited a significantly increased binding affinity than BA.2 (**Figure 3B**). Consistent with the results of pseudovirus assay (**Figure 3A**), these observations suggest that the N460K substitution is critical to characterize the virological phenotype of BA.2.75 S. To reveal the structural effect of the N460K substitution, we generated a structural model of BA.2.75 S RBD using AlphaFold2 (Mirdita et al., 2022). Calculating the electrostatic potential of this model in comparison with the S RBDs of B.1.1 and BA.2 showed that K460 of BA.2.75 S RBD is positively charged (**Figure 3C**), and the K460 is complementary to the negative charged binding site on human ACE2 (**Figure 3D**). These structural observations suggest that N460K substitution contributes to increased electrostatic complementary binding between the BA.2.75 S RBD and human ACE2.

Although the N460K substitution significantly increased binding affinity (**Figure 3B**), the binding affinity of the BA.2 N460K was still 5-fold lower than that of BA.2.75 (**Figure 3B**). Therefore, the extraordinary tight binding of BA.2.75 cannot be explained by the N460K alone, and it is hypothesized that the additional substitutions conferred negative effects in the BA.2 background. In particular, the D339H substitution requires two nucleotide changes in the codon to occur. Such changes are still relatively rare in the evolution of SARS-CoV-2, reinforcing the importance and corresponding fitness advantage. To analyze the potential impact of this substitution, we additionally prepared the BA.2.75 H339D derivative and measured its affinity. The K_D_ value of this mutant was significantly (3-fold) lower than that of the parental BA.2.75 (**Figure 3B**). The structural model computed by AlphaFold2 (Mirdita *et al*., 2022) suggested that the loss of ion-dipole interaction between the D339 and the N343 allowed for the N343 side chain repositioning (**Figure S3B**). These data suggest that the D339H substitution potentially influences the position of the linoleic acid binding loop between residues 367–378 (Toelzer et al., 2020) and thereby increases binding affinity to ACE2.

To further reveal the virological property of BA.2.75 S, we performed a cell-based fusion assay (Kimura et al., 2022b; Kimura *et al*., 2022c; Motozono et al., 2021; Saito *et al*., 2022; Suzuki *et al*., 2022; Yamasoba *et al*., 2022b) using Calu-3 cells as target cells. Flow cytometry analysis showed that the surface expression level of BA.2.75 is comparable to that of BA.2 (**Figure 3E**). Consistent with our recent study (Kimura *et al*., 2022c), the fusogenicity of BA.5 was significantly higher than that of BA.2, and notably, the BA.2.75 S was also significantly more fusogenic than the BA.2 S (**Figure 3F**). Altogether, these results suggest that BA.2.75 S exhibits higher binding affinity to human ACE2 and higher fusogenicity.

### Virological characteristics of BA.2.75 clinical isolate *in vitro*

To evaluate the growth capacity of BA.2.75, a clinical isolate of BA.2.75 (strain TY41-716; GISAID ID: EPI_ISL_13969765) was inoculated in a variety of *in vitro* cell culture systems. As controls, we also used clinical isolates of B.1.1 (strain TKYE610670; GISAID ID: EPI_ISL_479681) (Suzuki *et al*., 2022), Delta (B.1.617.2, strain TKYTK1734; GISAID ID: EPI_ISL_2378732) (Saito *et al*., 2022), BA.2 (strain TY40-385; GISAID ID: EPI_ISL_9595859) (Kimura *et al*., 2022c) and BA.5 (strain TKYS14631; GISAID ID: EPI_ISL_12812500) (Tamura *et al*., 2022). The growth efficacy of B.1.1 and Delta was significantly higher than that of BA.2 in Vero cells (**Figure 4A**), VeroE6/TMPRSS2 cells (**Figure 4B**), HEK293-ACE2/TMPRSS2 cells (**Figure 4C**), AO-derived air-liquid interface (AO-ALI) model (**Figure 4D**), human iPS cell (iPSC)-derived airway epithelial cells (**Figure 4E**) and lung epithelial cells (**Figure 4F**). BA.5 replicated more efficiently than BA.2 with statistically significant differences in the five cell culture systems except AO-ALI (**Figures 4A–4F**). The growth efficacy of BA.2.75 was significantly higher than that of BA.2 in Vero cells (**Figure 4A**), VeroE6/TMPRSS2 cells (**Figure 4B**), HEK293-ACE2/TMPRSS2 cells (**Figure 4C**), and iPSC-derived lung epithelial cells (**Figure 4F**), while the growth efficacy of BA.2.75 and BA.2 were comparable in the two airway epithelial cell systems (**Figures 4D and 4E**).

**Figure 4.**
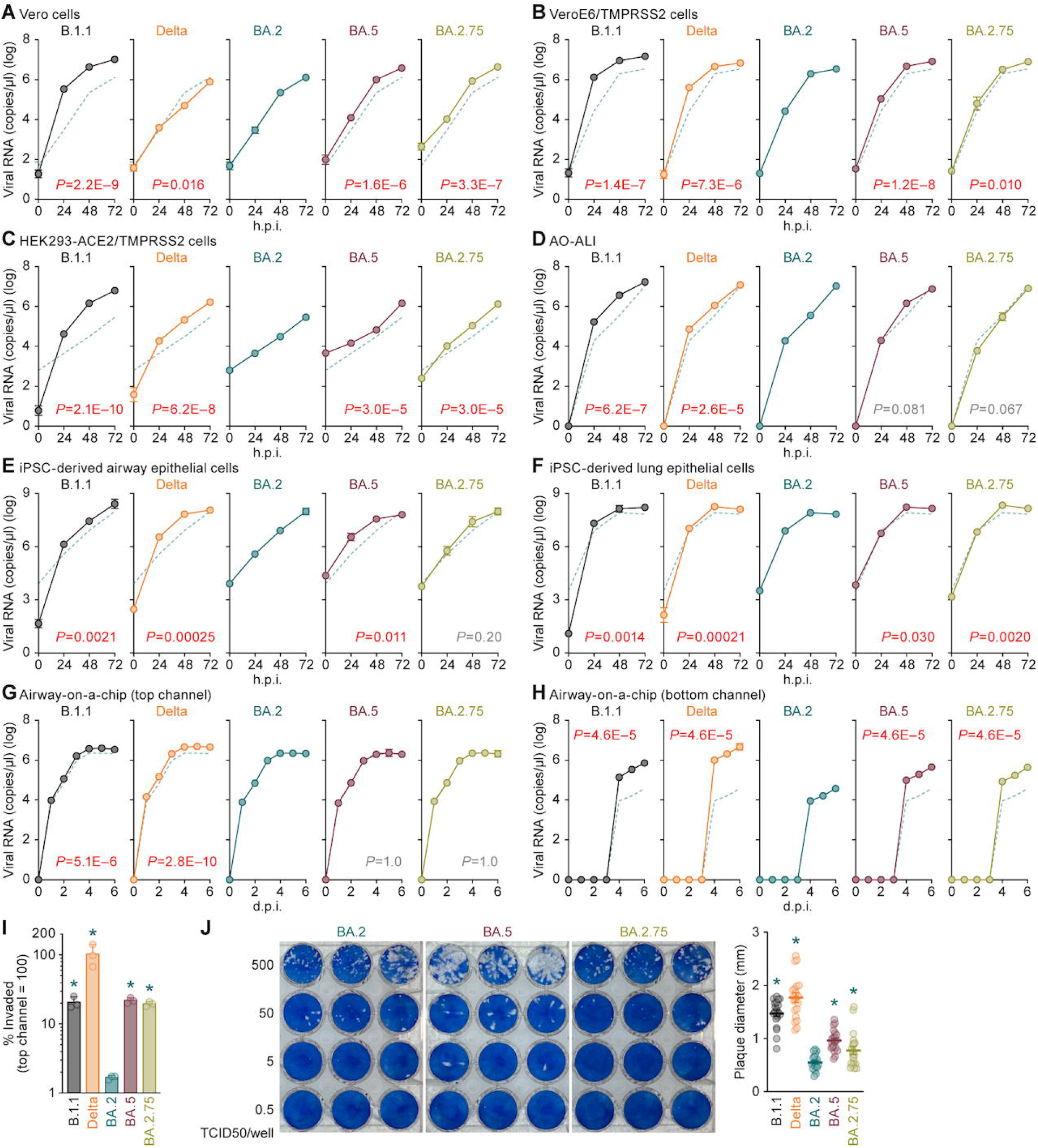
Growth capacity of BA.2.75 *in vitro*. (**A–I**) Growth kinetics of B.1.1, Delta, BA.2, BA.5 and BA.2.75. Clinical isolates of B.1.1 (strain TKYE610670; GISAID ID: EPI_ISL_479681), Delta (B.1.617.2, strain TKYTK1734; GISAID ID: EPI_ISL_2378732), BA.2 (strain TY40-385; GISAID ID: EPI_ISL_9595859), BA.5 (strain TKYS14631; GISAID ID: EPI_ISL_12812500), and BA.2.75 (strain TY41-716; GISAID ID: EPI_ISL_13969765) were inoculated into Vero cells (**A**), VeroE6/TMPRSS2 cells (**B**), HEK293-ACE2/TMPRSS2 cells (**C**), AO-ALI (**D**), iPSC-derived airway epithelial cells (**E**), iPSC-derived lung epithelial cells (**F**), and an airway-on-a-chip system (**G and H**; the scheme of experimental system is illustrated in Figure S3C). The copy numbers of viral RNA in the culture supernatant (**A–C**), the apical sides of cultures (**D–F**), the top (**G**) and bottom (**H**) channels of an airway-on-a-chip were routinely quantified by RT–qPCR. The dashed green line in each panel indicates the results of BA.2. In I, the percentage of viral RNA load in the bottom channel per top channel at 6 d.p.i. (i.e., % invaded virus from the top channel to the bottom channel) is shown. (**J**) Plaque assay. VeroE6/TMPRSS2 cells were used for the target cells. Representative panels (left) and a summary of the recorded plaque diameters (20 plaques per virus) (right) are shown. Assays were performed in quadruplicate, and the presented data are expressed as the average ± SD. In A–H, statistically significant differences between BA.2 and the other variants across timepoints were determined by multiple regression. The FWERs calculated using the Holm method are indicated in the figures. In I and J (right), statistically significant differences versus BA.2 (*, *P* < 0.05) were determined by two-sided Mann–Whitney *U* tests. Each dot indicates the result of an individual replicate. See also Figure S3.

To evaluate the effect of BA.2.75 on the airway epithelial and endothelial barriers, airway-on-a-chips (**Figure S3C**) were used. By measuring the amount of virus that invades from the top channel (airway channel; **Figure 4G**) to the bottom channel (blood vessel channel; **Figure 4H**), the ability of viruses to disrupt the airway epithelial and endothelial barriers can be evaluated. Notably, the amount of virus that invades to the blood vessel channel of BA.2.75-, BA.5- and B.1.1-infected airway-on-chips was significantly higher than that of BA.2-infected one (**Figure 4I**). These results suggest that BA.2.75 exhibits more severe airway epithelial and endothelial barrier disruption than BA.2.

To further address the fusogenic capacity of BA.2.75, we performed plaque assay using VeroE6/TMPRSS2 cells. Consistent with our previous studies using a Delta isolate (Saito *et al*., 2022) as well as the recombinant SARS-CoV-2 bearing the B.1.1 S (Yamasoba *et al*., 2022a), BA.2 S (Yamasoba *et al*., 2022a), and BA.5 S (Kimura *et al*., 2022c), the plaques formed by the infections of clinical isolates of B.1.1, Delta and BA.5 were significantly bigger than those formed by the infection of BA.2 (**Figure 4J**). Notably, BA.2.75 infection also showed significantly bigger plaques than BA.2 infection (**Figure 4J**). Together with the results of cell-based fusion assay (**Figure 3F**) and airway-on-a-chip infection experiments (**Figures 4G–4I**), these observations suggest that BA.2.75 is more fusogenic than BA.2, and the fusogenicity of BA.2.75 is comparable to that of BA.5.

### Virological characteristics of BA.2.75 *in vivo*

As we proposed in our prior studies (Kimura *et al*., 2022c; Saito *et al*., 2022; Suzuki *et al*., 2022; Yamasoba *et al*., 2022b), the fusogenicity of the S proteins of SARS-CoV-2 variants is closely associated with the intrinsic pathogenicity in an experimental hamster model. Here we revealed that both BA.5 and BA.2.75 are more fusogenic than BA.2 in the *in vitro* cell culture systems (**Figures 3 and 4**). Given that the recombinant SARS-CoV-2 bearing the BA.5 S (Kimura *et al*., 2022c) as well as a clinical isolate of BA.5 (Tamura *et al*., 2022) exhibited relatively higher pathogenicity than BA.2 in hamsters, it is hypothesized that BA.2.75 is also intrinsically more pathogenic than BA.2. To address this possibility, we intranasally inoculated a BA.2.75 isolate into hamsters. As controls, we also used clinical isolates of Delta, BA.2 and BA.5. While we followed our established experimental protocol (Kimura *et al*., 2022c; Saito *et al*., 2022; Suzuki *et al*., 2022; Yamasoba *et al*., 2022b), the viral titers of clinical isolates of Omicron subvariants were relatively low. Therefore, we set out to conduct animal experiments in this study with relatively lower titer inoculum (1,000 TCID_50_ per hamster) than our previous studies (10,000 TCID_50_ per hamster) (Kimura *et al*., 2022c; Saito *et al*., 2022; Suzuki *et al*., 2022; Yamasoba *et al*., 2022b). Nevertheless, consistent with our previous study (Saito *et al*., 2022), the Delta infection exhibited the most severe weight changes among the five groups (**Figure 5A**). While the body weight of BA.2-infected hamsters was similar to that of uninfected hamsters, those of BA.5- and BA.2.75-infected hamsters were significantly lower than that of uninfected hamsters (**Figure 5A**).

**Figure 5.**
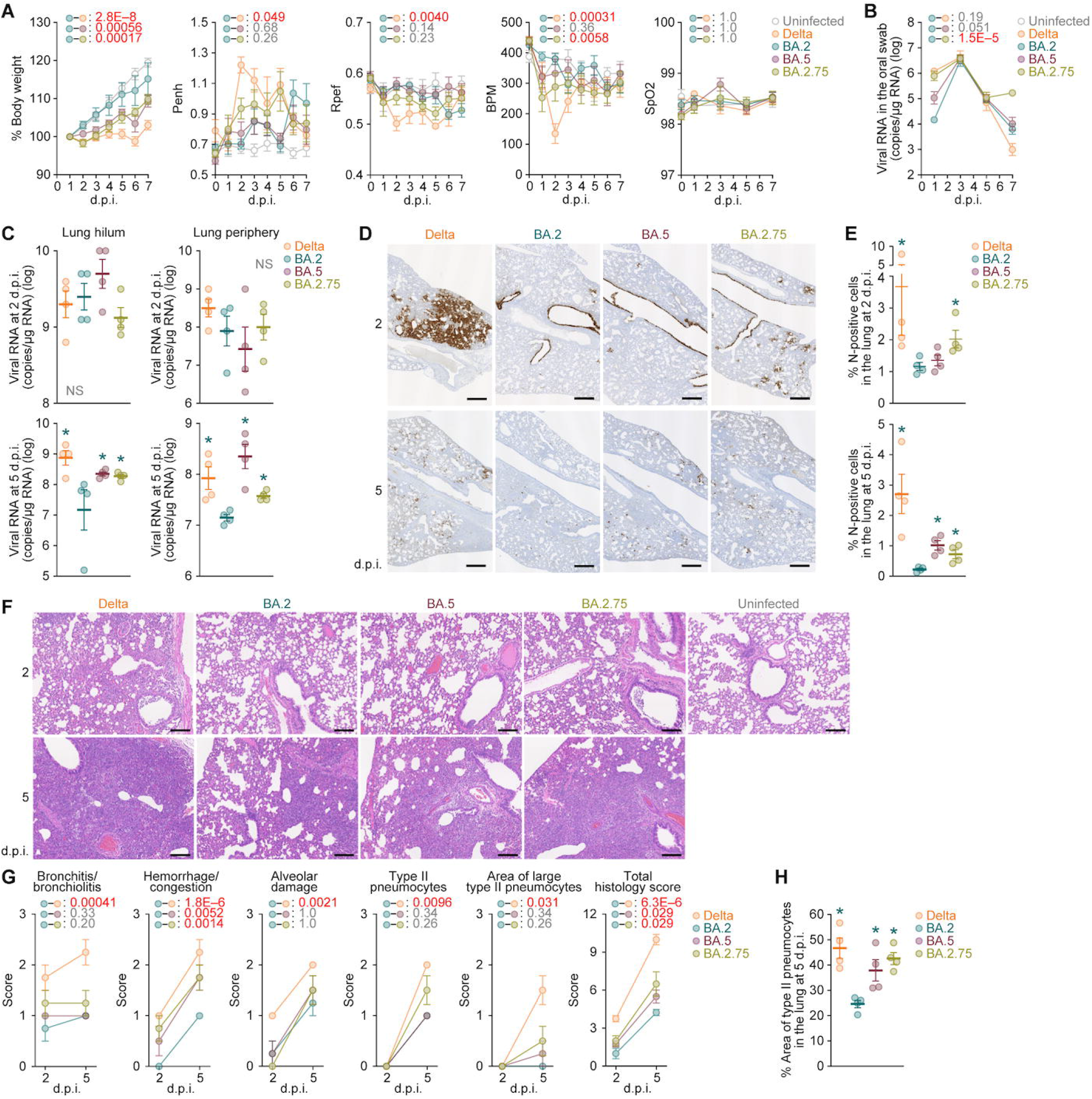
Virological characteristics of BA.2.75 *in vivo*. Syrian hamsters were intranasally inoculated with Delta, BA.2, BA.5 and BA.2.75. Six hamsters at the same age were intranasally inoculated with saline (uninfected). Six hamsters per each group were used to routinely measure respective parameters (A and B). Four hamsters per each group were euthanized at 2 and 5 d.p.i and used for virological and pathological analysis (**C–G**). (**A**) Body weight, Penh, Rpef, BPM, and SpO_2_ values of infected hamsters (n = 6 per infection group). (**B**) Viral RNA loads in the oral swab (n = 6 per infection group). (**C**) Viral RNA loads in the lung hilum (left) and lung periphery (right) of infected hamsters (n = 4 per infection group) at 2 d.p.i. (top) and 5 d.p.i. (bottom). (**D and E**) IHC of the viral N protein in the lungs at 2 d.p.i. (top) and 5 d.p.i. (bottom) of all infected hamsters. (**D**) Representative figures. (**E**) Percentage of N-positive cells in whole lung lobes (n = 4 per infection group). The raw data are shown in Figure S4B and S4C. (**F** and **G**) (**F**) H&E staining of the lungs of infected hamsters. Representative figures are shown. Uninfected lung alveolar space and bronchioles are also shown. (**G**) Histopathological scoring of lung lesions (n = 4 per infection group). Representative pathological features are reported in our previous studies (Kimura *et al*., 2022c; Saito *et al*., 2022; Suzuki *et al*., 2022; Yamasoba *et al*., 2022b). (H) Type II pneumocytes in the lungs of infected hamsters. The percentage of the area of type II pneumocytes in the lung at 5 d.p.i. is summarized. The raw data are shown in Figure S4D. In **A–C, E, G and H**, data are presented as the average ± SEM. In **C, E and H**, each dot indicates the result of an individual hamster. In **A, B and G**, statistically significant differences between BA.2 and other variants across timepoints were determined by multiple regression. In A, the 0 d.p.i. data were excluded from the analyses. The FWERs calculated using the Holm method are indicated in the figures. In **C, E and G**, the statistically significant differences between BA.2 and other variants were determined by a two-sided Mann–Whitney *U* test. In **D and F**, each panel shows a representative result from an individual infected hamster. Scale bars, 500 μm (D); 200 μm (F). See also Figure S4.

We then quantitatively analyzed the pulmonary function of infected hamsters as reflected by three parameters, enhanced pause (Penh), the ratio of time to peak expiratory follow relative to the total expiratory time (Rpef), and breath per minute (BPM), which are surrogate markers for bronchoconstriction or airway obstruction. Subcutaneous oxygen saturation (SpO_2_) was also routinely measured. Although the SpO_2_ values were comparable among the five groups, Delta infection resulted in significant differences in the other three respiratory parameters compared to BA.2 (**Figure 5A**), suggesting that Delta is more pathogenic than BA.2. There were no differences in the values of Penh, Rpef and BPM between BA.5 and BA.2, and the values of Penh and Rpef of BA.2.75-infected hamsters were comparable to those of BA.2 (**Figure 5A**). However, the BPM value of BA.2.75 was significantly lower than that of BA.2 (**Figure 5A**), suggesting that BA.2.75 is slightly more pathogenic than BA.2.

To address the viral spread in infected hamsters, we routinely measured the viral RNA load in the oral swab. Although the viral RNA loads of the hamsters infected with Delta, BA.2 and BA.5 were comparable, the viral load in the swabs of BA.2.75-infected hamsters was relatively highly maintained by 7 d.p.i. and was significantly higher than that of BA.2-infected hamsters (**Figure 5B**). To address the possibility that BA.2.75 more efficiently spread in the respiratory tissues, we collected the lungs of infected hamsters at 2 and 5 d.p.i., and the collected tissues were separated into the hilum and periphery regions. Although the viral RNA loads in both the hilum and periphery of four infection groups were comparable at 2 d.p.i. (**Figure 5C, top**), those of the hamsters infected with Delta, BA.5 and BA.2.75 were significantly higher than those infected with BA.2 at 5 d.p.i. (**Figure 5C, bottom**).

To further address the virus spread in the respiratory tissues, immunohistochemical (IHC) analysis targeting viral nucleocapsid (N) protein was conducted. Similar to our previous studies (Kimura *et al*., 2022c; Suzuki *et al*., 2022; Yamasoba *et al*., 2022b), epithelial cells in the upper tracheae of infected hamsters were sporadically positive for viral N protein at 2 d.p.i., but there were no significant differences among four viruses including BA.2.75 (**Figure S4A**). In the alveolar space around the bronchi/bronchioles at 2 d.p.i., the N-positive cells were detected in Delta-infected hamsters. On the other hand, the N proteins strongly remained in the lobar bronchi in BA.5- and BA.2.75-infected hamsters (**Figures 5D, top, and S4B**). While few N-positive cells were detected in the alveolar space of BA.2- and BA.5-infected hamsters, it was notable that the N positivity spread into the alveolar space in BA.2.75-infected hamsters (**Figures 5D, top, and S4B**). The quantification of the N-positive area in total of four lung lobes at 2 d.p.i. (**Figure S4B**) showed that the N-positive areas of Delta- and BA.2.75-infected hamsters were significantly greater than that of BA.2-infected hamsters (**Figure 5E, top**). At 5 d.p.i., although the N-positive cells were hardly detected in the lungs infected with BA.2, a few N-positive cells were detected in the peripheral alveolar space in Delta, BA.5, BA.2.75 (**Figures 5D, bottom, and S4C**). The quantification of the N-positive area in the four lung lobes at 5 d.p.i. (**Figure S4C**) further showed that the N-positive areas of Delta- and BA.5- and BA.2.75-infected hamsters were significantly greater than that of BA.2-infected hamsters (**Figure 5E, bottom**). These data suggest that BA.2 targets only a portion of bronchial/bronchiolar epithelium and was less efficiently transmitted to the neighboring epithelial cells. On the other hand, BA.5 and BA.2.75 infections seemed to persist in the bronchial/bronchiolar epithelium, and particularly, BA.2.75 invaded the alveolar space more efficiently than BA.5 at the early stage of infection. Altogether, the IHC data suggest that among Omicron subvariants, BA.2.75 more efficiently spread into the alveolar space than BA.2 and BA.5, with persistent infection in the bronchi/bronchioles.

### Pathogenicity of BA.2.75

To investigate the intrinsic pathogenicity of BA.2.75, the formalin-fixed right lungs of infected hamsters at 2 and 5 d.p.i. were analyzed by carefully identifying the four lobules and main bronchus and lobar bronchi sectioning each lobe along with the bronchial branches. Histopathological scoring was performed according to the criteria described in our previous studies (Kimura *et al*., 2022c; Saito *et al*., 2022; Suzuki *et al*., 2022; Yamasoba *et al*., 2022b): (i) bronchitis/bronchiolitis (an inflammatory indicator at early stage of infection), (ii) hemorrhage/congestion, (iii) alveolar damage with epithelial apoptosis and macrophage infiltration, (iv) emergence of type II pneumocytes, and (v) hyperplasia of type II pneumocytes were evaluated by certified pathologists and the degree of these pathological findings were arbitrarily scored using four-tiered system as 0 (negative), 1 (weak), 2 (moderate), and 3 (severe). Consistent with our previous studies (Saito *et al*., 2022; Suzuki *et al*., 2022), all five parameters as well as the total score of Delta-infected hamsters were significantly higher than those of BA.2-infected hamsters (**Figures 5F and 5G**), suggesting that Delta is more pathogenic than BA.2. When we compare the histopathological scores of Omicron subvariants, the scores indicating hemorrhage or congestion and total histology scores of BA.5 and BA.2.75 were significantly greater than those of BA.2 (**Figures 5F and 5G**). Similar to our recent studies (Kimura *et al*., 2022c; Tamura *et al*., 2022), BA.5 is intrinsically more pathogenic than BA.2, and notably, our results suggest that BA.2.75 exhibits more significant inflammation than BA.2. To clarify the area of pneumonia, the inflammatory area, which is mainly composed of the type II pneumocytes with some inflammatory cell types, such as neutrophils, lymphocytes, and macrophages, is termed the area of type II pneumocytes and was morphometrically analyzed (**Figure S4D**). As summarized in **Figure 5H**, at 5 d.p.i., the percentages of the area of type II pneumocytes of Delta, BA.5 and BA.2.75 were significantly higher than that of BA.2. Altogether, these findings suggest that BA.2.75 infection intrinsically induces greater inflammation and exhibits higher pathogenicity than BA.2.

## Discussion

Here, we characterized the virological property of the Omicron BA.2.75 variant, such as the growth rate in the human population, resistance to antiviral humoral immunity and antiviral drugs, functions of S protein *in vitro*, and intrinsic pathogenicity.

In terms of the emergence geography and phylogeny, BA.5 and BA.2.75 emerged independently. Nevertheless, the results of cell-based fusion assay, airway-on-a-chip assay and plaque assay suggested that both BA.5 and BA.2.75 acquired higher fusogenicity after the divergence from BA.2. Our data including a recent study (Kimura *et al*., 2022c) suggest that the critical substitution responsible for the higher fusogenicity of BA.5 and BA.2.75 S proteins are different: the L452R substitution for BA.5 S, and the D339H/N460K substitution for BA.2.75 S.

The higher fusogenicity attributed by the increased binding affinity of the L452R-bearing S RBD to human ACE2 was reported in previous studies focusing on the S proteins of previous SARS-CoV-2 variants including Epsilon (Motozono *et al*., 2021), Delta (Saito *et al*., 2022) and Omicron BA.5 (Kimura *et al*., 2022c) variants. The prominently increased ACE2 binding affinity caused by the N460K substitution was also reported in our previous study (Zahradnik et al., 2021b). We also demonstrated that the D339H, which is unique in the BA.2.75 S, contributes to increased ACE2 binding affinity. Our data suggest that the N460K and D339H substitutions cooperatively determine the higher fusogenicity of BA.2.75 S. In our previous studies focusing on Delta (Saito *et al*., 2022), Omicron BA.1 (Suzuki *et al*., 2022), BA.2 (Yamasoba *et al*., 2022b) and BA.5 (Kimura *et al*., 2022c), we proposed a close association between the S-mediated fusogenicity *in vitro* and the pathogenicity in a hamster model. Consistent with our hypothesis, here we demonstrated that, compared to BA.2, BA.2.75 exhibits higher fusogenicity *in vitro* and efficient viral spread in the lungs of infected hamsters, which leads to enhanced inflammation in the lung and higher pathogenicity *in vivo*. Moreover, *in vitro* experiments using a variety of cell culture systems showed that BA.2.75 replicates more efficiently than BA.2 in alveolar epithelial cells but not in airway epithelial cells. Altogether, our results suggest that BA.2.75 exhibits higher fusogenicity and pathogenicity via evolution of its S protein independently of BA.5.

Consistent with our previous study (Yamasoba *et al*., 2022b), neutralization experiments showed that BA.5 was significantly more resistant to the humoral immunity induced by vaccination and breakthrough infections of prior Omicron subvariants. On the other hand, the sensitivity of BA.2.75 to these antisera was comparable to BA.2. More importantly, BA.2.75 was highly resistant to the BA.5-induced immunity. These results suggest that, although both BA.2.75 and BA.5 are descendants of BA.2, their immunogenicity is different from each other. Furthermore, compared to BA.2, the sensitivity of BA.2.75 and BA.5 to therapeutic monoclonal antibodies was also different (Yamasoba *et al*., 2022a). The G446S was also closely associated with the resistance of BA.2.75 to the antiviral effects of BA.2- and BA.5-infected hamster sera. Because the G446S significantly decreases ACE2 affinity of S RBD, this substitution was acquired to evade antiviral immunity, and the other substitutions in RBD, particularly N460K, contributed to compensate for the decreased ACE2 binding affinity by G446S.

Another remarkable substitution pattern in the BA.2.75 S is the multiple substitutions in the S NTD. Particularly, three out of the five substitutions in the NTD (K147E, W152R and F157L) are located in a well-studied region, the NTD supersite. Previous studies showed that the mutations in the NTD supersite are responsible for the resistance to antiviral monoclonal antibodies (Cerutti *et al*., 2021; Chi *et al*., 2020; Liu *et al*., 2020; Lok, 2021; McCallum *et al*., 2021; Suryadevara *et al*., 2021; Voss *et al*., 2021). In fact, our results suggested that these three substitutions in the NTD supersite are closely associated with the evasion from BA.5-induced humoral immunity, in addition to the G446S in RBD. In fact, the W152 has been shown as a mutational hot spot of SARS-CoV-2 (Kubik et al., 2021). Therefore, BA.2.75 might mutate this specific residue to evade neutralization by sera of convalescent or vaccinated individuals.

According to the “COVID-19 Treatment Guidelines” issued by NIH (NIH, 2022), the use of Paxlovid (Ritonavir and Nirmatrelvir), Remdesivir and Molnupiravir (a prodrug of EIDD-1931) is highly recommended as treatment of patients who do not require hospitalization or oxygen supplement. Because the evolution of SARS-CoV-2 is unpredictable, timely and accurate testing of the efficacy of currently available antiviral drugs is indispensable to treat patients infected with a new variant. Our results using physiologically relevant human AO demonstrated that BA.2.75 retained the sensitivity to major small-molecule anti-SARS-CoV-2 drugs including Remdesivir, EIDD-1931 and Nirmatrelvir. Interestingly, BA.2.75 was more sensitive to Remdesivir than other stains, and a similar tendency was observed with EIDD-1931. In terms of drug testing, previous studies addressed the antiviral activity of these drugs against BA.2 and BA.5 using immortalized cell lines such as VeroE6/TMPRSS2 cells (Takashita et al., 2022c), Caco-2-F03 cells (Bojkova et al., 2022) and Calu-3 cells (Carlin et al., 2022) but the effects of these antiviral drugs were different each other, and these results were also different from ours (**Table 1**). In addition, a previous study demonstrated that VeroE6 cells have a low capacity to metabolize Remdesivir, leading to a weak antiviral activity (Pruijssers et al., 2020). These results suggest that the experimental system significantly affects the outcome of antiviral drug efficacy, raising the importance to evaluate the efficacy of antiviral drugs using physiologically relevant systems, such as organoids and organ-on-a-chip systems.

Our investigation using the viral genome surveillance data reported from India suggested that BA.2.75 bears the potential to outcompete BA.2 as well as BA.5, the most predominant variant in the world as of August 2022. Following the worldwide spread of BA.5, it is probable that the number of individuals infected with BA.5 will increase. Together with our findings showing the higher resistance of BA.2.75 to the BA.5-induced immunity, there appears to be sufficient plausibility that BA.2.75 evades the BA.5-induced immunity, and this property will confer this variant to more efficient spread in the countries where BA.5 has been widely spreading, such as Australia and Japan. Additionally, here we showed that the intrinsic pathogenicity of BA.2.75 in hamsters is comparable to BA.5 and higher than that of BA.2. Since a recent study showed that the hospitalization risk of BA.5 was significantly higher than that of BA.2 in the once-boosted vaccinated population (Kislaya et al., 2022), it is not unreasonable to infer that the intrinsic pathogenicity in infected hamsters reflects to the severity and outcome in infected humans to a meaningful extent.

In summary, our multiscale investigations revealed the growth rate in the human population, fusogenicity and intrinsic pathogenicity of BA.2.75 are greater than BA.2. These features of BA.2.75 suggests the potential risk of this variant to global health. Since BA.2.75 shows significantly higher R_e_ than BA.2 and BA.5 in India, this variant will probably transmit to and initiate outcompeting BA.2 and BA.5 in some countries other than India in the near future. To assess the potential risk of BA.2.75 to global health, this variant should be under monitoring carefully and continuously through worldwide cooperation of in-depth viral genomic surveillance.

## STAR*METHODS

● KEY RESOURCES TABLE
● RESOURCE AVAILABILITY

○ Lead Contact
○ Materials Availability
○ Data and Code Availability
○ EXPERIMENTAL MODEL AND SUBJECT DETAILS

○ Ethics Statement
○ Human serum collection
○ Cell culture
○ METHOD DETAILS

○ Viral genome sequencing
○ Phylogenetic analyses
○ Modelling the epidemic dynamics of SARS-CoV-2 lineages
○ Plasmid construction
○ Neutralization assay
○ Airway organoids
○ SARS-CoV-2 preparation and titration
○ Antiviral drug assay using SARS-CoV-2 clinical isolates and AO
○ Cytotoxicity assay
○ Pseudovirus infection
○ Yeast surface display
○ AlphaFold2
○ SARS-CoV-2 S-based fusion assay
○ AO-ALI model
○ Preparation of human airway and alveolar epithelial cells from human
○ iPSC

○ Airway-on-a-chips
○ Microfluidic device
○ SARS-CoV-2 infection
○ RT–qPCR
○ Plaque assay
○ Animal experiments
○ Lung function test
○ Immunohistochemistry
○ H&E staining
○ Histopathological scoring
○ QUANTIFICATION AND STATISTICAL ANALYSIS

## Supplemental Information

Additional Supplemental Items are available upon request.

## Author Contributions

Akatsuki Saito, Sayaka Deguchi, Izumi Kimura, Hesham Nasser, Mako Toyoda, Kayoko Nagata, Keiya Uriu, Yusuke Kosugi, Shigeru Fujita, Daichi Yamasoba, Maya Shofa, MST Monira Begum, Takashi Irie, Takamasa Ueno, and Terumasa Ikeda performed cell culture experiments.

Tomokazu Tamura, Koshiro Tabata, Rigel Suzuki, Hayato Ito, Naganori Nao, Kumiko Yoshimatsu, Hirofumi Sawa, Keita Matsuno, and Takasuke Fukuhara performed animal experiments.

Yoshitaka Oda, Lei Wang, Masumi Tsuda, and Shinya Tanaka performed histopathological analysis.

Jiri Zahradnik and Gideon Schreiber performed yeast surface display assay. Jiri Zahradnik and Yusuke Kosugi performed structural analysis.

Sayaka Deguchi and Kazuo Takayama prepared AO, AO-ALI and airway-on-a-chip systems.

Yuki Yamamoto and Tetsuharu Nagamoto performed generation and provision of human iPSC-derived airway and alveolar epithelial cells.

Hiroyuki Asakura, Mami Nagashima, Kenji Sadamasu, Kazuhisa Yoshimura performed viral genome sequencing analysis.

Akifumi Takaori-Kondo and Kotaro Shirakawa contributed clinical sample collection.

Jumpei Ito performed statistical, modelling, and bioinformatics analyses.

Jumpei Ito, Kazuo Takayama, Keita Matsuno, Shinya Tanaka, Terumasa Ikeda, Takasuke Fukuhara, and Kei Sato designed the experiments and interpreted the results.

Jumpei Ito, Terumasa Ikeda, Takasuke Fukuhara and Kei Sato wrote the original manuscript.

All authors reviewed and proofread the manuscript.

The Genotype to Phenotype Japan (G2P-Japan) Consortium contributed to the project administration.

## Acknowledgments

We would like to thank all members belonging to The Genotype to Phenotype Japan (G2P-Japan) Consortium. We thank Dr. Kenzo Tokunaga (National Institute for Infectious Diseases, Japan) and Dr. Jin Gohda (The University of Tokyo, Japan) for providing reagents. We also thank National Institute for Infectious Diseases, Japan for providing clinical isolates of BA.2.75 (strain TY41-716; GISAID ID: EPI_ISL_13969765) and BA.2 (strain TY40-385; GISAID ID: EPI_ISL_9595859) and Chiba University (Motoaki Seki, Ryoji Fujiki, Atsushi Kaneda, Tadanaga Shimada, Taka-aki Nakada, Seiichiro Sakao and Takuji Suzuki) for collecting and providing Delta infection sera. We gratefully acknowledge all data contributors, i.e. the Authors and their Originating laboratories responsible for obtaining the specimens, and their Submitting laboratories for generating the genetic sequence and metadata and sharing via the GISAID Initiative, on which this research is based. The super-computing resource was provided by Human Genome Center at The University of Tokyo.

This study was supported in part by AMED Program on R&D of new generation vaccine including new modality application (JP223fa727002, to Kei Sato); AMED Research Program on Emerging and Re-emerging Infectious Diseases (JP21fk0108574, to Hesham Nasser; JP21fk0108465, to Akatsuki Saito; JP21fk0108493, to Takasuke Fukuhara; JP22fk0108617 to Takasuke Fukuhara; JP22fk0108146, to Kei Sato; 21fk0108494 to G2P-Japan Consortium, Kotaro Shirakawa, Takashi Irie, Keita Matsuno, Shinya Tanaka, Terumasa Ikeda, Takasuke Fukuhara, and Kei Sato); AMED Research Program on HIV/AIDS (JP22fk0410033, to Akatsuki Saito; JP22fk0410047, to Akatsuki Saito; JP22fk0410055, to Terumasa Ikeda; 22fk0410034 to Akifumi Takaori-Kondo and Kotaro Shirakawa; and JP22fk0410039, to Kotaro Shirakawa and Kei Sato); AMED CRDF Global Grant (JP22jk0210039 to Akatsuki Saito); AMED Japan Program for Infectious Diseases Research and Infrastructure (JP22wm0325009, to Akatsuki Saito; JP22wm0125008 to Keita Matsuno); AMED CREST (JP22gm1610005, to Kazuo Takayama); JST CREST (JPMJCR20H4, to Kei Sato); JSPS KAKENHI Grant-in-Aid for Scientific Research C (22K07089, to Mako Toyoda; 22K07103, to Terumasa Ikeda); JSPS KAKENHI Grant-in-Aid for Scientific Research B (21H02736, to Takasuke Fukuhara); JSPS KAKENHI Grant-in-Aid for Early-Career Scientists (22K16375, to Hesham Nasser; 20K15767, Jumpei Ito); JSPS Core-to-Core Program (A. Advanced Research Networks) (JPJSCCA20190008, to Kei Sato); JSPS Research Fellow DC2 (22J11578, to Keiya Uriu); JSPS Leading Initiative for Excellent Young Researchers (LEADER) (to Terumasa Ikeda); World-leading Innovative and Smart Education (WISE) Program 1801 from the Ministry of Education, Culture, Sports, Science and Technology (MEXT) (to Naganori Nao); The Tokyo Biochemical Research Foundation (to Kei Sato); Takeda Science Foundation (to Terumasa Ikeda); Shin-Nihon Foundation of Advanced Medical Research (to Mako Toyoda and Terumasa Ikeda); Waksman Foundation of Japan (to Terumasa Ikeda); an intramural grant from Kumamoto University COVID-19 Research Projects (AMABIE) (to Terumasa Ikeda).

## Consortia

Mai Kishimoto, Marie Kato, Zannatul Ferdous, Hiromi Mouri, Kenji Shishido, Naoko Misawa, Mai Suganami, Mika Chiba, Ryo Yoshimura, So Nakagawa, Jiaqi Wu, Yasuhiro Kazuma, Ryosuke Nomura, Yoshihito Horisawa, Yusuke Tashiro, Yugo Kawai, Ryoko Kawabata, Ryo Shimizu, Otowa Takahashi, Kimiko Ichihara, Chihiro Motozono, Yuri L. Tanaka, Erika P. Butlertanaka, Rina Hashimoto, Takao Hashiguchi, Tateki Suzuki, Kanako Kimura, Jiei Sasaki, Yukari Nakajima, Kaori Tabata

**Table S1.** Estimated relative R_e_ values of viral lineages in India, related to **Figure 1**

**Table S2.** Human sera used in this study, related to **Figure 2**

**Table S3.** Primers used for the construction of SARS-CoV-2 S expression plasmids, related to **Figures 2 and 3**

**Table S4.** Summary of unexpected amino acid mutations detected in the working virus stocks, related to **Figures 4 and 5 and** Table 1

**Figure S1. Epidemic dynamics of BA.2.75 in India, related to Figure 1**

(**A**) Amino acid differences in B.1.1, Delta, BA.2, BA.5 and BA.2.75 compared to the SARS-CoV-2 A lineage. Heatmap color indicates the frequency of amino acid mutations.

(**B**) Estimated relative R_e_ of each viral lineage, assuming a fixed generation time of 2.1 days. The R_e_ value of BA.2 is set at 1. The posterior (violin), posterior mean (dot), and 95% CI (line) are shown. The R_e_ values for respective Indian states are shown. The dynamics of the top seven predominant lineages in India were estimated. BA.5 sublineages are summarized as “BA.5”, and non-predominant BA.2 sublineages are summarized as “other BA.2”. Raw data are summarized in **Table S1**.

(**C**) Fold change in R_e_ values between BA.2.75 and BA.5. Posterior mean (dot) and 95% CI (line) are shown. Red indicates that the 95% CI does not overlap with the value of 1.

**Figure S2. Effects of antiviral drugs in AO, related to Table 1**

(A) Antiviral effects of the three drugs in AO culture. The assay of each antiviral drugs was performed in quadruplicate, and the 50% effective concentration (EC_50_) was calculated. The data are summarized in **Table 1**.

(B) Cytotoxic effects of the three drugs in AO culture. The assay of each antiviral drugs was performed in quadruplicate, and the 50% cytotoxic concentration (CC_50_) was calculated. The CC_50_ values are indicated in the panels.

**Figure S3. Virological features of BA.2.75 *in vitro*, related to Figures 3 and 4**

(A) Fold increase in pseudovirus infectivity based on TMPRSS2 expression.

(B) The structural effect of the D339H substitution in the BA.2.75 S RBD. The BA.2 S RBD (PDB: 7UB0) (Stalls *et al*., 2022) and an AlphaFold2 structural model of BA.2.75 S RBD (bottom) are shown. The residues 339 and 343 are indicated in stick. The squared regions are enlarged in the right panel. A dashed line in the top panel indicates ion-dipole interaction between the D339 and the N343 residues.

(C) A scheme of airway-on-a-chip system.

**Figure S4. Histological observations in infected hamsters, related to Figure 5**

(**A**) IHC of the viral N protein in the middle portion of the tracheas of all infected hamsters at 2 d.p.i (4 hamsters per infection group). Each panel shows a representative result from an individual infected hamster.

(**B and C**) IHC of the SARS-CoV-2 N protein in the lungs of infected hamsters at 2 d.p.i. (**B**) and 5 d.p.i (**C**) (4 hamsters per infection group). In each panel, IHC staining (top) and the digitalized N-positive area (bottom, indicated in red) are shown. The red numbers in the bottom panels indicate the percentage of the N-positive area. Summarized data are shown in Figure 5E.

(D) Type II pneumocytes in the lungs of infected hamsters (4 hamsters per infection group). H&E staining (top) and the digitalized inflammatory area with type II pneumocytes (bottom, indicated in red) are shown. The red numbers in the bottom panels indicate the percentage of inflammatory area with type II pneumocytes. Summarized data are shown in Figure 5H.

Scale bars, 1 mm (**A**); 5 mm (**B–D**).

## STAR*METHODS

### KEY RESOURCES TABLE

### RESOURCE AVAILABILITY

#### Lead Contact

Further information and requests for resources and reagents should be directed to and will be fulfilled by the Lead Contact, Kei Sato.

#### Materials Availability

All unique reagents generated in this study are listed in the Key Resources Table and available from the Lead Contact with a completed Materials Transfer Agreement.

#### Data and Software Availability

All databases/datasets used in this study are available from GISAID database (https://www.gisaid.org) and GenBank database (https://www.gisaid.org; EPI_SET ID: EPI_SET_220804hy).

The computational codes used in the present study, the raw data of virus sequences, and the GISAID supplemental table for EPI_SET ID: EPI_SET_220804hy are available in the GitHub repository (https://github.com/TheSatoLab/Omicron_BA.2.75).

### EXPERIMENTAL MODEL AND SUBJECT DETAILS

#### Ethics statement

All experiments with hamsters were performed in accordance with the Science Council of Japan’s Guidelines for the Proper Conduct of Animal Experiments. The protocols were approved by the Institutional Animal Care and Use Committee of National University Corporation Hokkaido University (approval ID: 20-0123 and 20-0060). All experiments with mice were also performed in accordance with the Science Council of Japan’s Guidelines for the Proper Conduct of Animal Experiments. All protocols involving specimens from human subjects recruited at Kyoto University were reviewed and approved by the Institutional Review Boards of Kyoto University (approval ID: G1309) and Chiba University (approval ID: HS202103-03). All human subjects provided written informed consent. All protocols for the use of human specimens were reviewed and approved by the Institutional Review Boards of The Institute of Medical Science, The University of Tokyo (approval IDs: 2021-1-0416 and 2021-18-0617), Kyoto University (approval ID: G0697), Kumamoto University (approval IDs: 2066 and 2074), and University of Miyazaki (approval ID: O-1021).

#### Human serum collection

Vaccine sera of fifteen individuals who had BNT162b2 vaccine (Pfizer/BioNTech) (average age: 38, range: 24–48; 53% male) (**Figures 2A–2C**) were obtained at one month after the second dose, one month after the third dose, and four months after the third dose. The details of the vaccine sera are summarized in **Table S2**.

Convalescent sera were collected from the following donors: fully vaccinated individuals who had been infected with BA.1 (16 2-dose vaccinated. 10–27 days after testing; average age: 48, range: 20–76, 44% male) (**Figure 2D**), fully vaccinated individuals who had been infected with BA.2 (9 2-dose vaccinated and 5 3-dose vaccinated. 11–61 days after testing. n=14 in total; average age: 47, range: 24–84, 64% male) (**Figure 2E**), and unvaccinated individuals who had been infected with Delta (6–55 days after testing. n=18 in total; average age: 50, range: 22–67, 78% male) (**Figure 2F**). The SARS-CoV-2 variants were identified as previously described (Kimura *et al*., 2022c; Yamasoba *et al*., 2022b). Sera were inactivated at 56°C for 30 minutes and stored at –80°C until use. The details of the convalescent sera are summarized in **Table S2**.

#### Cell culture

HEK293T cells (a human embryonic kidney cell line; ATCC, CRL-3216), HEK293 cells (a human embryonic kidney cell line; ATCC, CRL-1573) and HOS-ACE2/TMPRSS2 cells (HOS cells stably expressing human ACE2 and TMPRSS2) (Ferreira et al., 2021; Ozono et al., 2021) were maintained in DMEM (high glucose) (Sigma-Aldrich, Cat# 6429-500ML) containing 10% fetal bovine serum (FBS, Sigma-Aldrich Cat# 172012-500ML) and 1% penicillin-streptomycin (PS) (Sigma-Aldrich, Cat# P4333-100ML).

HEK293-ACE2 cells (HEK293 cells stably expressing human ACE2) (Motozono *et al*., 2021) were maintained in DMEM (high glucose) containing 10% FBS, 1 µg/ml puromycin (InvivoGen, Cat# ant-pr-1) and 1% PS.

HEK293-ACE2/TMPRSS2 cells (HEK293 cells stably expressing human ACE2 and TMPRSS2) (Motozono *et al*., 2021) were maintained in DMEM (high glucose) containing 10% FBS, 1 µg/ml puromycin, 200 ng/ml hygromycin (Nacalai Tesque, Cat# 09287-84) and 1% PS.

Vero cells [an African green monkey (*Chlorocebus sabaeus*) kidney cell line; JCRB Cell Bank, JCRB0111] were maintained in Eagle’s minimum essential medium (EMEM) (Sigma-Aldrich, Cat# M4655-500ML) containing 10% FBS and 1% PS.

VeroE6/TMPRSS2 cells (VeroE6 cells stably expressing human TMPRSS2; JCRB Cell Bank, JCRB1819) (Matsuyama et al., 2020) were maintained in DMEM (low glucose) (Wako, Cat# 041-29775) containing 10% FBS, G418 (1 mg/ml; Nacalai Tesque, Cat# G8168-10ML) and 1% PS.

Calu-3/DSP_1-7_ cells (Calu-3 cells stably expressing DSP_1-7_) (Yamamoto et al., 2020) were maintained in EMEM (Wako, Cat# 056-08385) containing 20% FBS and 1% PS.

Human airway and alveolar epithelial cells derived from human induced pluripotent stem cells (iPSCs) were manufactured according to established protocols as described below (see “Preparation of human airway and alveolar epithelial cells from human iPSCs” section) and provided by HiLung Inc.

Airway organoids (AO) and AO-derived air-liquid interface model (AO-ALI) were generated according to established protocols as described below (see “Airway organoids” and “AO-ALI model” sections).

### METHOD DETAILS

#### Viral genome sequencing

Viral genome sequencing was performed as previously described (Meng *et al*., 2022; Motozono *et al*., 2021; Saito *et al*., 2022; Suzuki *et al*., 2022; Yamasoba *et al*., 2022b). Briefly, the virus sequences were verified by viral RNA-sequencing analysis. Viral RNA was extracted using a QIAamp viral RNA mini kit (Qiagen, Cat# 52906). The sequencing library employed for total RNA sequencing was prepared using the NEB next ultra RNA library prep kit for Illumina (New England Biolabs, Cat# E7530). Paired-end 76-bp sequencing was performed using a MiSeq system (Illumina) with MiSeq reagent kit v3 (Illumina, Cat# MS-102-3001). Sequencing reads were trimmed using fastp v0.21.0 (Chen et al., 2018) and subsequently mapped to the viral genome sequences of a lineage B isolate (strain Wuhan-Hu-1; GenBank accession number: NC_045512.2) (Matsuyama *et al*., 2020) using BWA-MEM v0.7.17 (Li and Durbin, 2009). Variant calling, filtering, and annotation were performed using SAMtools v1.9 (Li et al., 2009) and snpEff v5.0e (Cingolani et al., 2012).

#### Phylogenetic analyses

To construct an ML tree of Omicron lineages (BA.1–BA.5) sampled from South Africa and BA.2.75 (shown in **Figure 1A**), the genome sequence data of SARS-CoV-2 and its metadata were downloaded from the GISAID database (https://www.gisaid.org/) (Khare et al., 2021) on July 23, 2022. We excluded the data of viral strains with the following features from the analysis: i) a lack collection date information; ii) sampling from animals other than humans, iii) >2% undetermined nucleotide characters, or iv) sampling by quarantine. From each viral lineage, 30 sequences were randomly sampled and used for tree construction, in addition to an outgroup sequence, EPI_ISL_466615, representing the oldest isolate of B.1.1 obtained in the UK. The viral genome sequences were mapped to the reference sequence of Wuhan-Hu-1 (GenBank accession number: NC_045512.2) using Minimap2 v2.17 (Li, 2018) and subsequently converted to a multiple sequence alignment according to the GISAID phylogenetic analysis pipeline (https://github.com/roblanf/sarscov2phylo). The alignment sites corresponding to the 1–265 and 29674–29903 positions in the reference genome were masked (i.e., converted to NNN). Alignment sites at which >50% of sequences contained a gap or undetermined/ambiguous nucleotide were trimmed using trimAl v1.2 (Capella-Gutierrez et al., 2009). Phylogenetic tree construction was performed via a three-step protocol: i) the first tree was constructed; ii) tips with longer external branches (Z score > 4) were removed from the dataset; iii) and the final tree was constructed. Tree reconstruction was performed by RAxML v8.2.12 (Stamatakis, 2014) under the GTRCAT substitution model. The node support value was calculated by 100 times bootstrap analysis.

#### Modelling the epidemic dynamics of SARS-CoV-2 lineages

To quantify the spread rate of each SARS-CoV-2 lineage in the human population in India, we estimated the relative R_e_ of each viral lineage according to the epidemic dynamics, calculated on the basis of viral genomic surveillance data. The data were downloaded from the GISAID database (https://www.gisaid.org/) on August 1, 2022. We excluded the data of viral strains with the following features from the analysis: i) a lack of collection date information; ii) sampling in animals other than humans; or iii) sampling by quarantine. We analyzed the datasets of the ten states of India, where ≥20 sequences of either BA.2.75 or BA.5 are reported (i.e., Himachal Pradesh, Odisha, Haryana, Rajasthan, and Maharashtra, Gujarat, West Bengal, Delhi, Tamil Nadu, and Telangana). BA.5 sublineages are summarized as “BA.5”, and BA.2 sublineages with ≤400 sequences are summarized as “other BA.2”. Subsequently, the dynamics of the top seven predominant lineages in India were estimated from April 24, 2022, to August 1, 2022, were analyzed. The number of viral sequences of each viral lineage collected on each day in each country was counted, and the count matrix was constructed as an input for the statistical model below.

We constructed a Bayesian hierarchical model to represent relative lineage growth dynamics with multinomial logistic regression as described in our previous study (Yamasoba *et al*., 2022b). In brief, we incorporated a hierarchical structure into the slope parameter over time, which enabled us to estimate the global average relative R_e_ of each viral lineage in India as well as the average value for each country. Arrays in the model index over one or more indices: L = 7 viral lineages *l*; S = 10 states *s*; and T = 100 days *t*; The model is:

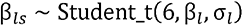

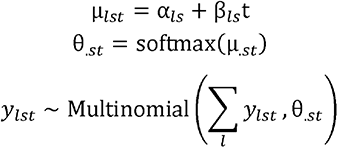

The explanatory variable was time, *t*, and the outcome variable was *y_lst_*, which represented the count of viral lineage *l* in state *s* at time *t*. The slope parameter of lineage *l* in state *s*, β*_ls_*, was generated from a Student’s *t* distribution with hyperparameters of the mean, β*_l_*, and the standard deviation, *σ_l_*. As the distribution generating *β_ls_*, we used a Student’s t distribution with six degrees of freedom instead of a normal distribution to reduce the effects of outlier values of β*_ls_*. In the model, the linear estimator µ_.*st*_, consisting of the intercept α_.*s*_ and the slope β_.*s*_, was converted to the simplex θ*_.st_*, which represented the probability of occurrence of each viral lineage at time *t* in state *s*, based on the softmax link function defined as:

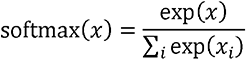

*y_lst_* is generated from θ*_.st_* and the total count of all lineages at time *t* in state *s* according to a multinomial distribution.

The relative R_e_ of each viral lineage in each county (*r_ls_*) was calculated according to the slope parameter β*_ls_* as:

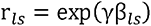

Where γ is the average viral generation time (2.1 days) (http://sonorouschocolate.com/covid19/index.php?title=Estimating_Generation_Time_Of_Omicron). Similarly, the global average relative R_e_ of each viral lineage was calculated according to the slope hyperparameter β*_l_* as:

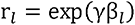

For parameter estimation, the intercept and slope parameters of the BA.2 variant were fixed at 0. Consequently, the relative R_e_ of BA.2 was fixed at 1, and those of the other lineages were estimated relative to that of BA.2.

Parameter estimation was performed via the MCMC approach implemented in CmdStan v2.28.1 (https://mc-stan.org) with CmdStanr v0.4.0 (https://mc-stan.org/cmdstanr/). Noninformative priors were set for all parameters. Four independent MCMC chains were run with 1,000 and 2,000 steps in the warmup and sampling iterations, respectively. We confirmed that all estimated parameters showed <1.01 R-hat convergence diagnostic values and >200 effective sampling size values, indicating that the MCMC runs were successfully convergent. The above analyses were performed in R v4.1.3 (https://www.r-project.org/). Information on the relative R_e_ estimated in the present study is summarized in **Table S1**.

#### Plasmid construction

Plasmids expressing the codon-optimized SARS-CoV-2 S proteins of B.1.1 (the parental D614G-bearing variant), BA.2 and BA.5 were prepared in our previous studies (Kimura et al., 2022a; Ozono *et al*., 2021; Saito *et al*., 2022; Suzuki *et al*., 2022; Yamasoba *et al*., 2022b). Plasmids expressing the codon-optimized S proteins of BA.2.75 and the BA.2 S-based derivatives were generated by site-directed overlap extension PCR using the primers listed in **Table S3**. The resulting PCR fragment was digested with KpnI and NotI and inserted into the corresponding site of the pCAGGS vector (Niwa et al., 1991). Nucleotide sequences were determined by DNA sequencing services (Eurofins), and the sequence data were analyzed by Sequencher v5.1 software (Gene Codes Corporation).

#### Neutralization assay

Pseudoviruses were prepared as previously described (Kimura *et al*., 2022a; Kimura *et al*., 2022b; Kimura *et al*., 2022c; Motozono *et al*., 2021; Uriu et al., 2022; Uriu et al., 2021; Yamasoba *et al*., 2022a; Yamasoba *et al*., 2022b; Yamasoba *et al*., 2022c). Briefly, lentivirus (HIV-1)-based, luciferase-expressing reporter viruses were pseudotyped with the SARS-CoV-2 S proteins. HEK293T cells (1,000,000 cells) were cotransfected with 1 μg psPAX2-IN/HiBiT (Ozono et al., 2020), 1 μg pWPI-Luc2 (Ozono *et al*., 2020), and 500 ng plasmids expressing parental S or its derivatives using PEI Max (Polysciences, Cat# 24765-1) according to the manufacturer’s protocol. Two days posttransfection, the culture supernatants were harvested and centrifuged. The pseudoviruses were stored at –80°C until use.

Neutralization assay (**Figure 2**) was prepared as previously described (Kimura *et al*., 2022a; Kimura *et al*., 2022b; Kimura *et al*., 2022c; Saito *et al*., 2022; Uriu *et al*., 2022; Uriu *et al*., 2021; Yamasoba *et al*., 2022a; Yamasoba *et al*., 2022b; Yamasoba *et al*., 2022c). Briefly, the SARS-CoV-2 S pseudoviruses (counting ∼20,000 relative light units) were incubated with serially diluted (120-fold to 87,480-fold dilution at the final concentration) heat-inactivated sera at 37°C for 1 hour. Pseudoviruses without sera were included as controls. Then, a 40 μl mixture of pseudovirus and serum/antibody was added to HOS-ACE2/TMPRSS2 cells (10,000 cells/50 μl) in a 96-well white plate. At 2 d.p.i., the infected cells were lysed with a One-Glo luciferase assay system (Promega, Cat# E6130) or a Bright-Glo luciferase assay system (Promega, Cat# E2650), and the luminescent signal was measured using a GloMax explorer multimode microplate reader 3500 (Promega) or CentroXS3 (Berthhold Technologies). The assay of each serum was performed in triplicate, and the 50% neutralization titer (NT_50_) was calculated using Prism 9 software v9.1.1 (GraphPad Software).

#### Airway organoids

Airway organoids (AO) model was generated according to our previous report (Sano *et al*., 2022). Briefly, normal human bronchial epithelial cells (NHBE, Cat# CC-2540, Lonza) were used to generate AO. NHBE were suspended in 10 mg/ml cold Matrigel growth factor reduced basement membrane matrix (Corning). 50 μl of cell suspension was solidified on pre-warmed cell-culture treated multi-dishes (24-well plates; Thermo Fisher Scientific) at 37 °C for 10 min, and then 500 μl of expansion medium was added to each well. AO were cultured with AO expansion medium for 10 days. To mature the AO, expanded AO were cultured with AO differentiation medium for 5 days. In experiments evaluating the antiviral drugs (see “Antiviral drug assay using SARS-CoV-2 clinical isolates and AO” section below), AO were dissociated into single cells, and then were seeded into 96-well plates.

#### SARS-CoV-2 preparation and titration

The working virus stocks of SARS-CoV-2 were prepared and titrated as previously described (Kimura *et al*., 2022b; Kimura *et al*., 2022c; Meng *et al*., 2022; Motozono *et al*., 2021; Saito *et al*., 2022; Suzuki *et al*., 2022; Yamasoba *et al*., 2022b). In this study, clinical isolates of B.1.1 (strain TKYE610670; GISAID ID: EPI_ISL_479681) (Suzuki *et al*., 2022), Delta (B.1.617.2, strain TKYTK1734; GISAID ID: EPI_ISL_2378732) (Saito *et al*., 2022), BA.2 (strain TY40-385; GISAID ID: EPI_ISL_9595859) (Kimura *et al*., 2022c) and BA.5 (strain TKYS14631; GISAID ID: EPI_ISL_12812500) (Tamura *et al*., 2022), and BA.2.75 (strain TY41-716; GISAID ID: EPI_ISL_13969765) were used. In brief, 20 μl of the seed virus was inoculated into VeroE6/TMPRSS2 cells (5,000,000 cells in a T-75 flask). One h.p.i., the culture medium was replaced with DMEM (low glucose) (Wako, Cat# 041-29775) containing 2% FBS and 1% PS. At 3 d.p.i., the culture medium was harvested and centrifuged, and the supernatants were collected as the working virus stock.

The titer of the prepared working virus was measured as the 50% tissue culture infectious dose (TCID_50_). Briefly, one day before infection, VeroE6/TMPRSS2 cells (10,000 cells) were seeded into a 96-well plate. Serially diluted virus stocks were inoculated into the cells and incubated at 37°C for 4 days. The cells were observed under microscopy to judge the CPE appearance. The value of TCID_50_/ml was calculated with the Reed–Muench method (Reed and Muench, 1938).

To verify the sequences of SARS-CoV-2 working viruses, viral RNA was extracted from the working viruses using a QIAamp viral RNA mini kit (Qiagen, Cat# 52906) and viral genome sequences were analyzed as described above (see “Viral genome sequencing” section). Information on the unexpected substitutions detected is summarized in **Table S4**, and the raw data are deposited in the GitHub repository (https://github.com/TheSatoLab/Omicron_BA.2.75).

#### Antiviral drug assay using SARS-CoV-2 clinical isolates and AO

Antiviral drug assay (**Table 1 and Figure S2A**) was performed as previously described (Meng *et al*., 2022). Briefly, one day before infection, AO (10,000 cells) was dissociated, and then seeded into a 96-well plate. The cells were infected with either B.1.1, BA.2, BA.5 or BA.2.75 isolate (100 TCID_50_) at 37□°C for 2 hours. The cells were washed with DMEM and cultured in DMEM supplemented with 10%□FCS, 1% PS and the serially diluted Remdesivir (Clinisciences, Cat# A17170), EIDD-1931 (an active metabolite of Molnupiravir; Cell Signalling Technology, Cat# 81178S), or Nirmatrelvir (PF-07321332; MedChemExpress, Cat# HY-138687). At 24□hours after infection, the culture supernatants were collected, and viral RNA was quantified using RT–qPCR (see “RT–qPCR” section below). The assay of each compound was performed in quadruplicate, and the 50% effective concentration (EC_50_) was calculated using Prism 9 software v9.1.1 (GraphPad Software).

#### Cytotoxicity assay

The cytotoxicity of Remdesivir, EIDD-1931 or Nirmatrelvir (**Figure S2B**) was performed as previously described (Meng *et al*., 2022). Briefly, one day before the assay, AO (10,000 cells) was dissociated and then seeded into a 96-well plate. The cells were cultured with the serially diluted antiviral drugs for 24□hours. The cell counting kit-8 (Dojindo, Cat# CK04-11) solution (10□ µl) was added to each well, and the cells were incubated at 37□°C for 90□min. Absorbance was measured at 450 nm using the Multiskan FC (Thermo Fisher Scientific). The assay of each compound was performed in quadruplicate, and the 50% cytotoxic concentration (CC_50_) was calculated using Prism 9 software v9.1.1 (GraphPad Software).

#### Pseudovirus infection

Pseudovirus infection (**Figure 3A**) was performed as previously described (Ferreira *et al*., 2021; Kimura *et al*., 2022a; Kimura *et al*., 2022b; Kimura *et al*., 2022c; Motozono *et al*., 2021; Uriu *et al*., 2022; Uriu *et al*., 2021; Yamasoba *et al*., 2022a; Yamasoba *et al*., 2022b; Yamasoba *et al*., 2022c). Briefly, the amount of pseudoviruses prepared was quantified by the HiBiT assay using Nano Glo HiBiT lytic detection system (Promega,Cat# N3040) as previously described (Ozono *et al*., 2021; Ozono *et al*., 2020), and the same amount of pseudoviruses (normalized to the HiBiT value, which indicates the amount of p24 HIV-1 antigen) was inoculated into HOS-ACE2/TMPRSS2 cells, HEK293-ACE2 cells or HEK293-ACE2/TMPRSS2 and viral infectivity was measured as described above (see “Neutralization assay” section). To analyze the effect of TMPRSS2 for pseudovirus infectivity (**Figure S3A**), the fold change of the values of HEK293-ACE2/TMPRSS2 to HEK293-ACE2 was calculated.

#### Yeast surface display

Yeast surface display (**Figure 3B**) was performed as previously described (Dejnirattisai *et al*., 2022; Kimura *et al*., 2022a; Kimura *et al*., 2022b; Kimura *et al*., 2022c; Motozono *et al*., 2021; Yamasoba *et al*., 2022b; Zahradnik *et al*., 2021b). Briefly, the RBD genes [“construct 3” in (Zahradnik *et al*., 2021b), covering residues 330–528] in pJYDC1 plasmid were cloned by restriction enzyme-free cloning and transformed into the EBY100 Saccharomyces cerevisiae. Primers are listed in **Table S3**. The expression media 1/9 (Zahradnik *et al*., 2021a) was inoculated (OD 1) by overnight (220 rpm, 30°C, SD-CAA media) grown culture and cultivated for 24 hours at 20°C. The media was supplemented by 10 mM DMSO solubilized bilirubin (Sigma-Aldrich, Cat# 14370-1G) for expression co-cultivation labeling (pJYDC1, eUnaG2 reporter holo-form formation, green/yellow fluorescence (Ex. 498 nm, Em. 527 nm). Cells (100 ul aliquots) were collected by centrifugation (3000 g, 3 minutes), washed in ice-cold PBSB buffer (PBS with 1 g/L BSA), and resuspended in an analysis solution with a series of CF®640R succinimidyl ester labeled (Biotium, USA, Cat# 92108) ACE2 peptidase domain (residues 18–740) concentrations. The reaction volume was adjusted (1–100 ml) to avoid the ligand depletion effect, and the suspension was incubated overnight in a rotator shaker (10 rpm, 4°C). Incubated samples were washed by PBSB buffer, transferred into 96 well plates (Thermo, USA, Nunc, Cat# 268200), and analyzed by the CytoFLEX S Flow Cytometer (Beckman Coulter, USA, Cat#. N0-V4-B2-Y4) with the gating strategy described previously (Zahradnik *et al*., 2021b). The eUnaG2 signals were compensated by the instrument CytExpert software (Beckman Coulter). The mean binding signal (FL4-A) values of RBD expressing cells, subtracted by signals of non-expressing populations, were subjected for the determination of binding constant KD, YD by non-cooperative Hill equation fitted by nonlinear least-squares regression using Python v3.7 fitted together with two additional parameters describing titration curve (Zahradnik *et al*., 2021b).

#### AlphaFold2

To generate the structure model of BA.2.75 S RBD (**Figures 3C, 3D and S3B**), the AlphaFold2 structural prediction was performed using ColabFold (Mirdita *et al*., 2022) using the BA.2 S RBD template (PDB: 7UB0) (Stalls *et al*., 2022). The MMseqs2 and HHsearch parameters were set as default. The models were manually inspected, and those exhibiting poor parameters and models that did not adopt the classical RBD interface conformation were eliminated. The two highest score models were analyzed in detail. Three-dimensional visualization and analyses were performed using PyMOL v2.1.1 (Schrödinger, https://pymol.org/2/). In **Figure 3D**, the wild-type SARS-CoV-2 S RBD of the crystal structure of RBD-ACE2 complex (PDB:6M17) (Yan *et al*., 2020) was replaced with the BA.2.75 S RBD structure generated by AlphaFold2.

#### SARS-CoV-2 S-based fusion assay

SARS-CoV-2 S-based fusion assay (**Figures 3E and 3F**) was performed as previously described (Kimura *et al*., 2022b; Kimura *et al*., 2022c; Motozono *et al*., 2021; Saito *et al*., 2022; Suzuki *et al*., 2022; Yamasoba *et al*., 2022b). Briefly, on day 1, effector cells (i.e., S-expressing cells) and target cells (Calu-3/DSP_1-7_ cells) were prepared at a density of 0.6–0.8 × 10^6^ cells in a 6-well plate. On day 2, to prepare effector cells, HEK293 cells were cotransfected with the S expression plasmids (400 ng) and pDSP_8-11_ (Kondo et al., 2011) (400 ng) using TransIT-LT1 (Takara, Cat# MIR2300). On day 3 (24 hours posttransfection), 16,000 effector cells were detached and reseeded into 96-well black plates (PerkinElmer, Cat# 6005225), and target cells were reseeded at a density of 1,000,000 cells/2 ml/well in 6-well plates. On day 4 (48 hours posttransfection), target cells were incubated with EnduRen live cell substrate (Promega, Cat# E6481) for 3 hours and then detached, and 32,000 target cells were added to a 96-well plate with effector cells. *Renilla* luciferase activity was measured at the indicated time points using Centro XS3 LB960 (Berthhold Technologies). To measure the surface expression level of S protein, effector cells were stained with rabbit anti-SARS-CoV-2 S S1/S2 polyclonal antibody (Thermo Fisher Scientific, Cat# PA5-112048, 1:100). Normal rabbit IgG (SouthernBiotech, Cat# 0111-01, 1:100) was used as negative controls, and APC-conjugated goat anti-rabbit IgG polyclonal antibody (Jackson ImmunoResearch, Cat# 111-136-144, 1:50) was used as a secondary antibody. Surface expression level of S proteins (**Figure 3E**) was measured using FACS Canto II (BD Biosciences) and the data were analyzed using FlowJo software v10.7.1 (BD Biosciences). To calculate fusion activity, *Renilla* luciferase activity was normalized to the MFI of surface S proteins. The normalized value (i.e., *Renilla* luciferase activity per the surface S MFI) is shown as fusion activity.

#### AO-ALI model

AO-ALI model (**Figure 4D**) was generated according to our previous report (Sano *et al*., 2022). To generate AO-ALI, expanding AO were dissociated into single cells, and then were seeded into Transwell inserts (Corning) in a 24-well plate. To promote their maturation, AO-ALI were cultured with AO differentiation medium for 5 days. AO-ALI were infected with SARS-CoV-2 from the apical side.

#### Preparation of human airway and alveolar epithelial cells from human Ipsc

The air-liquid interface culture of airway and alveolar epithelial cells (**Figures 4E and 4F**) were differentiated from human iPSC-derived lung progenitor cells as previously described (Gotoh et al., 2014; Kimura *et al*., 2022c; Konishi et al., 2016; Tamura *et al*., 2022; Yamamoto et al., 2017). Briefly, lung progenitor cells were stepwise induced from human iPSCs referring a 21-days and 4-steps protocol (Yamamoto *et al*., 2017). At day 21, lung progenitor cells were isolated with specific surface antigen carboxypeptidase M and seeded onto upper chamber of 24-well Cell Culture Insert (Falcon, #353104), followed by 28-day and 7-day differentiation of airway and alveolar epithelial cells, respectively. Alveolar differentiation medium supplemented with dexamethasone (Sigma-Aldrich, Cat# D4902), KGF (PeproTech, Cat# 100-19), 8-Br-cAMP (Biolog, Cat# B007), 3-Isobutyl 1-methylxanthine (IBMX) (FUJIFILM Wako, Cat# 095-03413), CHIR99021 (Axon Medchem, Cat# 1386), and SB431542 (FUJIFILM Wako, Cat# 198-16543) was used for induction of alveolar epithelial cells. PneumaCult ALI (STEMCELL Technologies, Cat# ST-05001) supplemented with heparin (Nacalai Tesque, Cat# 17513-96) and Y-27632 (LC Laboratories, Cat# Y-5301) hydrocortisone (Sigma-Aldrich, Cat# H0135) was used for induction of airway epithelial cells.

#### Airway-on-a-chips

Airway-on-a-chips (**Figure S3C**) were prepared as previously described (Hashimoto et al., 2022). Human lung microvascular endothelial cells (HMVEC-L) were obtained from Lonza (Cat# CC-2527) and cultured with EGM-2-MV medium (Lonza, Cat# CC-3202). To prepare the airway-on-a-chip, first, the bottom channel of a polydimethylsiloxane (PDMS) device was pre-coated with fibronectin (3 μg/ml, Sigma, Cat# F1141). The microfluidic device was generated according to our previous report (Deguchi et al., 2021). HMVEC-L were suspended at 5,000,000 cells/ml in EGM2-MV medium. Then, 10 μl suspension medium was injected into the fibronectin-coated bottom channel of the PDMS device. Then, the PDMS device was turned upside down and incubated for 1 h. After 1 hour, the device was turned over, and the EGM2-MV medium was added into the bottom channel. After 4 days, AO were dissociated and seeded into the top channel. The AO was generated according to our previous report (Sano *et al*., 2022). AO were dissociated into single cells and then suspended at 5,000,000 cells/ml in the AO differentiation medium. Ten microliter suspension medium was injected into the top channel. After 1 hour, the AO differentiation medium was added to the top channel. In the infection experiments (**Figures 4G–4I**), the AO differentiation medium containing either B.1.1, Delta, BA.2, BA.5 or BA.2.75 isolate (500 TCID_50_) was inoculated from the top channel (**Figure S3C**). At 2 h.p.i., the top and bottom channels were washed and cultured with AO differentiation and EGM2-MV medium, respectively. The culture supernatants were collected, and viral RNA was quantified using RT–qPCR (see “RT–qPCR” section above).

#### Microfluidic device

The microfluidic device was generated according to our previous report (Deguchi *et al*., 2021). Briefly, the microfluidic device consisted of two layers of microchannels separated by a semipermeable membrane. The microchannel layers were fabricated from PDMS using a soft lithographic method. PDMS prepolymer (SYLGARD 184, Dow Corning) at a base to curing agent ratio of 10:1 was cast against a mold composed of SU-8 2150 (MicroChem) patterns formed on a silicon wafer. The cross-sectional size of the microchannels was 1 mm in width and 330 μm in height. To introduce solutions into the microchannels, access holes were punched through the PDMS using a 6-mm biopsy punch (Kai Corporation). Two PDMS layers were bonded to a PET membrane containing 3.0 μm pores (Cat# 353091, Falcon) using a thin layer of liquid PDMS prepolymer as the mortar. PDMS prepolymer was spin-coated (4000 rpm for 60 sec) onto a glass slide. Subsequently, both the top and bottom channel layers were placed on the glass slide to transfer the thin layer of PDMS prepolymer onto the embossed PDMS surfaces. The membrane was then placed onto the bottom layer and sandwiched with the top layer. The combined layers were left at room temperature for 1 day to remove air bubbles and then placed in an oven at 60°C overnight to cure the PDMS glue. The PDMS devices were sterilized by placing them under UV light for 1 hour before the cell culture.

#### SARS-CoV-2 infection

One day before infection, Vero cells (10,000 cells), VeroE6/TMPRSS2 cells (10,000 cells), and HEK293-ACE2/TMPRSS2 cells were seeded into a 96-well plate. SARS-CoV-2 [1,000 TCID_50_ for Vero cells (**Figure 4A**); 100 TCID_50_ for VeroE6/TMPRSS2 cells (**Figure 4B**) and HEK293-ACE2/TMPRSS2 cells (**Figure 4C**)] was inoculated and incubated at 37°C for 1 hour. The infected cells were washed, and 180 µl culture medium was added. The culture supernatant (10 µl) was harvested at the indicated timepoints and used for RT–qPCR to quantify the viral RNA copy number (see “RT–qPCR” section below). In the infection experiments using human iPSC-derived airway and alveolar epithelial cells (**Figures 4E and 4F**), working viruses were diluted with Opti-MEM (Thermo Fisher Scientific, 11058021). The diluted viruses (1,000 TCID_50_ in 100□μl) were inoculated onto the apical side of the culture and incubated at 37□°C for 1□hour. The inoculated viruses were removed and washed twice with Opti-MEM. To collect the viruses, 100□μl Opti-MEM was applied onto the apical side of the culture and incubated at 37□°C for 10□minutes. The Opti-MEM was collected and used for RT–qPCR to quantify the viral RNA copy number (see “RT–qPCR” section below). The infection experiments using an airway-on-a-chip system (**Figures 4G–4I**) was performed as described above (see “Airway-on-a-chips” section).

#### RT–qPCR

RT–qPCR was performed as previously described (Kimura *et al*., 2022b; Kimura *et al*., 2022c; Meng *et al*., 2022; Motozono *et al*., 2021; Saito *et al*., 2022; Suzuki *et al*., 2022; Yamasoba *et al*., 2022b). Briefly, 5 μl culture supernatant was mixed with 5 μl 2 × RNA lysis buffer [2% Triton X-100 (Nacalai Tesque, Cat# 35501-15), 50 mM KCl, 100 mM Tris-HCl (pH 7.4), 40% glycerol, 0.8 U/μl recombinant RNase inhibitor (Takara, Cat# 2313B)] and incubated at room temperature for 10 min. RNase-free water (90 μl) was added, and the diluted sample (2.5 μl) was used as the template for real-time RT-PCR performed according to the manufacturer’s protocol using One Step TB Green PrimeScript PLUS RT-PCR kit (Takara, Cat# RR096A) and the following primers: Forward *N*, 5’-AGC CTC TTC TCG TTC CTC ATC AC-3’; and Reverse *N*, 5’-CCG CCA TTG CCA GCC ATT C-3’. The viral RNA copy number was standardized with a SARS-CoV-2 direct detection RT-qPCR kit (Takara, Cat# RC300A). Fluorescent signals were acquired using QuantStudio 1 Real-Time PCR system (Thermo Fisher Scientific), QuantStudio 3 Real-Time PCR system (Thermo Fisher Scientific), QuantStudio 5 Real-Time PCR system (Thermo Fisher Scientific), CFX Connect Real-Time PCR Detection system (Bio-Rad), Eco Real-Time PCR System (Illumina), qTOWER3 G Real-Time System (Analytik Jena) Thermal Cycler Dice Real Time System III (Takara) or 7500 Real-Time PCR System (Thermo Fisher Scientific).

#### Plaque assay

Plaque assay (**Figure 4J**) was performed as previously described (Kimura *et al*., 2022b; Kimura *et al*., 2022c; Suzuki *et al*., 2022; Yamasoba *et al*., 2022b). Briefly, one day before infection, VeroE6/TMPRSS2 cells (100,000 cells) were seeded into a 24-well plate and infected with SARS-CoV-2 (0.5, 5, 50 and 500 TCID_50_) at 37°C for 1 hour. Mounting solution containing 3% FBS and 1.5% carboxymethyl cellulose (Wako, Cat# 039-01335) was overlaid, followed by incubation at 37°C. At 3 d.p.i., the culture medium was removed, and the cells were washed with PBS three times and fixed with 4% paraformaldehyde phosphate (Nacalai Tesque, Cat# 09154-85). The fixed cells were washed with tap water, dried, and stained with staining solution [0.1% methylene blue (Nacalai Tesque, Cat# 22412-14) in water] for 30 minutes. The stained cells were washed with tap water and dried, and the size of plaques was measured using Fiji software v2.2.0 (ImageJ).

#### Animal experiments

Animal experiments (**Figure 5**) were performed as previously described (Kimura *et al*., 2022b; Kimura *et al*., 2022c; Saito *et al*., 2022; Suzuki *et al*., 2022; Tamura *et al*., 2022; Yamasoba *et al*., 2022b). Syrian hamsters (male, 4 weeks old) were purchased from Japan SLC Inc. (Shizuoka, Japan). For the virus infection experiments, hamsters were euthanized by intramuscular injection of a mixture of 0.15 mg/kg medetomidine hydrochloride (Domitor^®^, Nippon Zenyaku Kogyo), 2.0 mg/kg midazolam (Dormicum^®^, FUJIFILM Wako Chemicals) and 2.5 mg/kg butorphanol (Vetorphale^®^, Meiji Seika Pharma) or 0.15 mg/kg medetomidine hydrochloride, 4.0 mg/kg alphaxaone (Alfaxan^®^, Jurox) and 2.5 mg/kg butorphanol. The Delta, BA.2, BA.5 and BA.2.75 (1,000 TCID_50_ in 100 µl), or saline (100 µl) were intranasally inoculated under anesthesia. Oral swabs were collected at indicated timepoints. Body weight was recorded daily by 7 d.p.i. Enhanced pause (Penh), the ratio of time to peak expiratory follow relative to the total expiratory time (Rpef), and BPM were measured every day until 7 d.p.i. (see below). Subcutaneous oxygen saturation (SpO_2_, see below) was monitored at 0, 1, 3, 5, and 7 d.p.i. Lung tissues were anatomically collected at 2 and 5 d.p.i. Viral RNA load in the oral swabs and respiratory tissues were determined by RT–qPCR. These tissues were also used for IHC and histopathological analyses (see below).

#### Lung function test

Lung function test (**Figure 5A**) was routinely performed as previously described (Kimura *et al*., 2022b; Kimura *et al*., 2022c; Saito *et al*., 2022; Suzuki *et al*., 2022; Tamura *et al*., 2022; Yamasoba *et al*., 2022b). The three respiratory parameters (Penh, Rpef and BPM) were measured by using a whole-body plethysmography system (DSI) according to the manufacturer’s instructions. In brief, a hamster was placed in an unrestrained plethysmography chamber and allowed to acclimatize for 30 seconds, then, data were acquired over a 2.5-minute period by using FinePointe Station and Review softwares v2.9.2.12849 (STARR). The state of oxygenation was examined by measuring SpO_2_ using pulse oximeter, MouseOx PLUS (STARR). SpO_2_ was measured by attaching a measuring chip to the neck of hamsters sedated by 0.25 mg/kg medetomidine hydrochloride.

#### Immunohistochemistry

Immunohistochemistry (IHC) (**Figures 5D, S4A–S4C**) was performed as previously described (Kimura *et al*., 2022b; Kimura *et al*., 2022c; Saito *et al*., 2022; Suzuki *et al*., 2022; Tamura *et al*., 2022; Yamasoba *et al*., 2022b) using an Autostainer Link 48 (Dako). The deparaffinized sections were exposed to EnVision FLEX target retrieval solution high pH (Agilent, Cat# K8004) for 20 minutes at 97°C to activate, and mouse anti-SARS-CoV-2 N monoclonal antibody (clone 1035111, R&D systems, Cat# MAB10474-SP, 1:400) was used as a primary antibody. The sections were sensitized using EnVision FLEX (Agilent) for 15 minutes and visualized by peroxidase-based enzymatic reaction with 3,3’-diaminobenzidine tetrahydrochloride (Dako, Cat# DM827) as substrate for 5 minutes. The N protein positivity (**Figures 5E, S4A and S4B**) was evaluated by certificated pathologists as previously described (Kimura *et al*., 2022b; Kimura *et al*., 2022c; Saito *et al*., 2022; Suzuki *et al*., 2022; Tamura *et al*., 2022; Yamasoba *et al*., 2022b). Images were incorporated as virtual slide by NDP.scan software v3.2.4 (Hamamatsu Photonics). The N-protein positivity was measured as the area using Fiji software v2.2.0 (ImageJ).

#### H&E staining

H&E staining (**Figures 5F** **and S4D**) was performed as previously described (Kimura *et al*., 2022b; Kimura *et al*., 2022c; Saito *et al*., 2022; Suzuki *et al*., 2022; Tamura *et al*., 2022; Yamasoba *et al*., 2022b). Briefly, excised animal tissues were fixed with 10% formalin neutral buffer solution, and processed for paraffin embedding. The paraffin blocks were sectioned with 3 µm-thickness and then mounted on MAS-GP-coated glass slides (Matsunami Glass, Cat# S9901). H&E staining was performed according to a standard protocol.

#### Histopathological scoring

Histopathological scoring (**Figure 5G**) was performed as previously described (Kimura *et al*., 2022b; Kimura *et al*., 2022c; Saito *et al*., 2022; Suzuki *et al*., 2022; Tamura *et al*., 2022; Yamasoba *et al*., 2022b). Pathological features including (i) bronchitis or bronchiolitis, (ii) hemorrhage with congestive edema, (iii) alveolar damage with epithelial apoptosis and macrophage infiltration, (iv) hyperplasia of type II pneumocytes, and (v) the area of the hyperplasia of large type II pneumocytes were evaluated by certified pathologists and the degree of these pathological findings were arbitrarily scored using four-tiered system as 0 (negative), 1 (weak), 2 (moderate), and 3 (severe). The “large type II pneumocytes” are the hyperplasia of type II pneumocytes exhibiting more than 10-μm-diameter nucleus. We described “large type II pneumocytes” as one of the remarkable histopathological features reacting SARS-CoV-2 infection in our previous studies (Kimura *et al*., 2022b; Kimura *et al*., 2022c; Saito *et al*., 2022; Suzuki *et al*., 2022; Tamura *et al*., 2022; Yamasoba *et al*., 2022b). Total histology score is the sum of these five indices.

To measure the inflammation area in the infected lungs (**Figures 5H** **and S4D**), four hamsters infected with each virus were sacrificed at 5 d.p.i., and all four right lung lobes, including upper (anterior/cranial), middle, lower (posterior/caudal), and accessory lobes, were sectioned along with their bronchi. The tissue sections were stained by H&E, and the digital microscopic images were incorporated into virtual slides using NDP.scan software v3.2.4 (Hamamatsu Photonics). The inflammatory area including type II pneumocyte hyperplasia in the infected whole lungs was morphometrically analyzed using Fiji software v2.2.0 (ImageJ).

### QUANTIFICATION AND STATISTICAL ANALYSIS

Statistical significance was tested using a two-sided Mann–Whitney *U*-test, a two-sided Student’s *t*-test or a two-sided paired *t-*test unless otherwise noted. The tests above were performed using Prism 9 software v9.1.1 (GraphPad Software).

In the time-course experiments (**Figures 3F, 4A–4H, 5A, 5B, and 5G**), a multiple regression analysis including experimental conditions (i.e., the types of infected viruses) as explanatory variables and timepoints as qualitative control variables was performed to evaluate the difference between experimental conditions thorough all timepoints. The initial time point was removed from the analysis. *P* value was calculated by a two-sided Wald test. Subsequently, familywise error rates (FWERs) were calculated by the Holm method. These analyses were performed in R v4.1.2 (https://www.r-project.org/).

In **Figures 5D, 5F and S4**, photographs shown are the representative areas of at least two independent experiments by using four hamsters at each timepoint.

