## Supplementary figures and images for "Virological characteristics of the SARS-CoV-2 Omicron BA.2.75"

### Figure S1

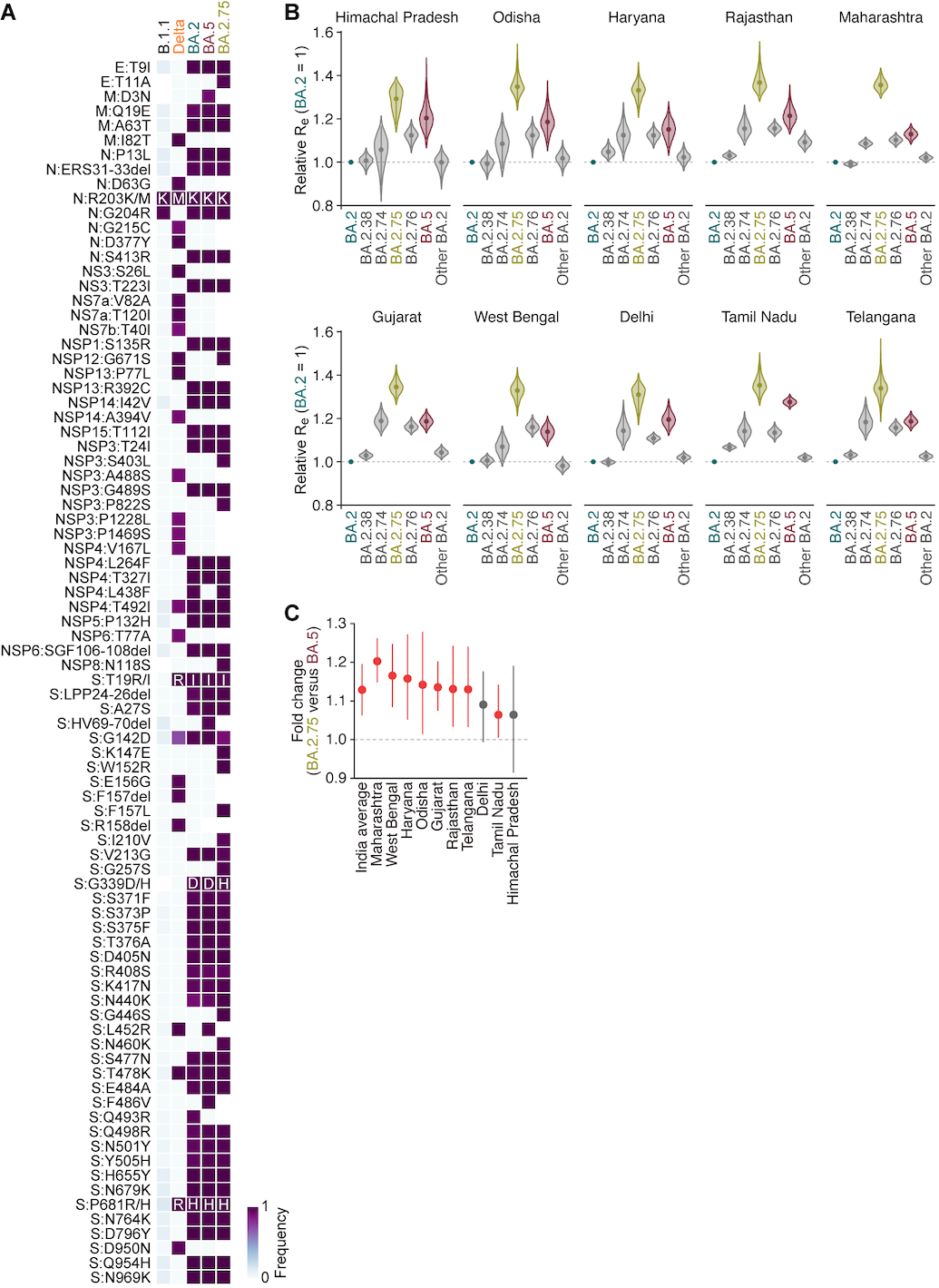

### Figure S2

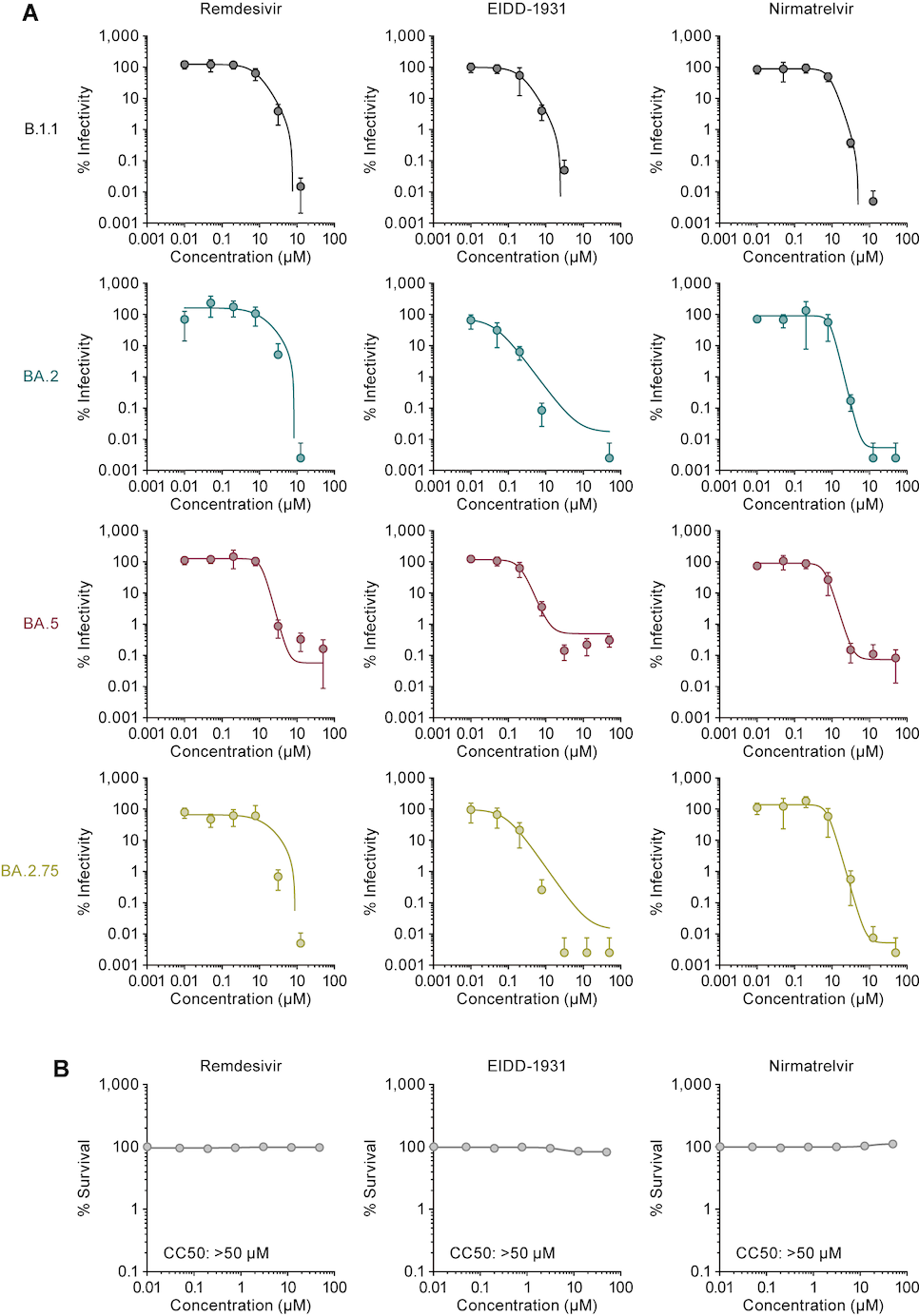

### Figure S3

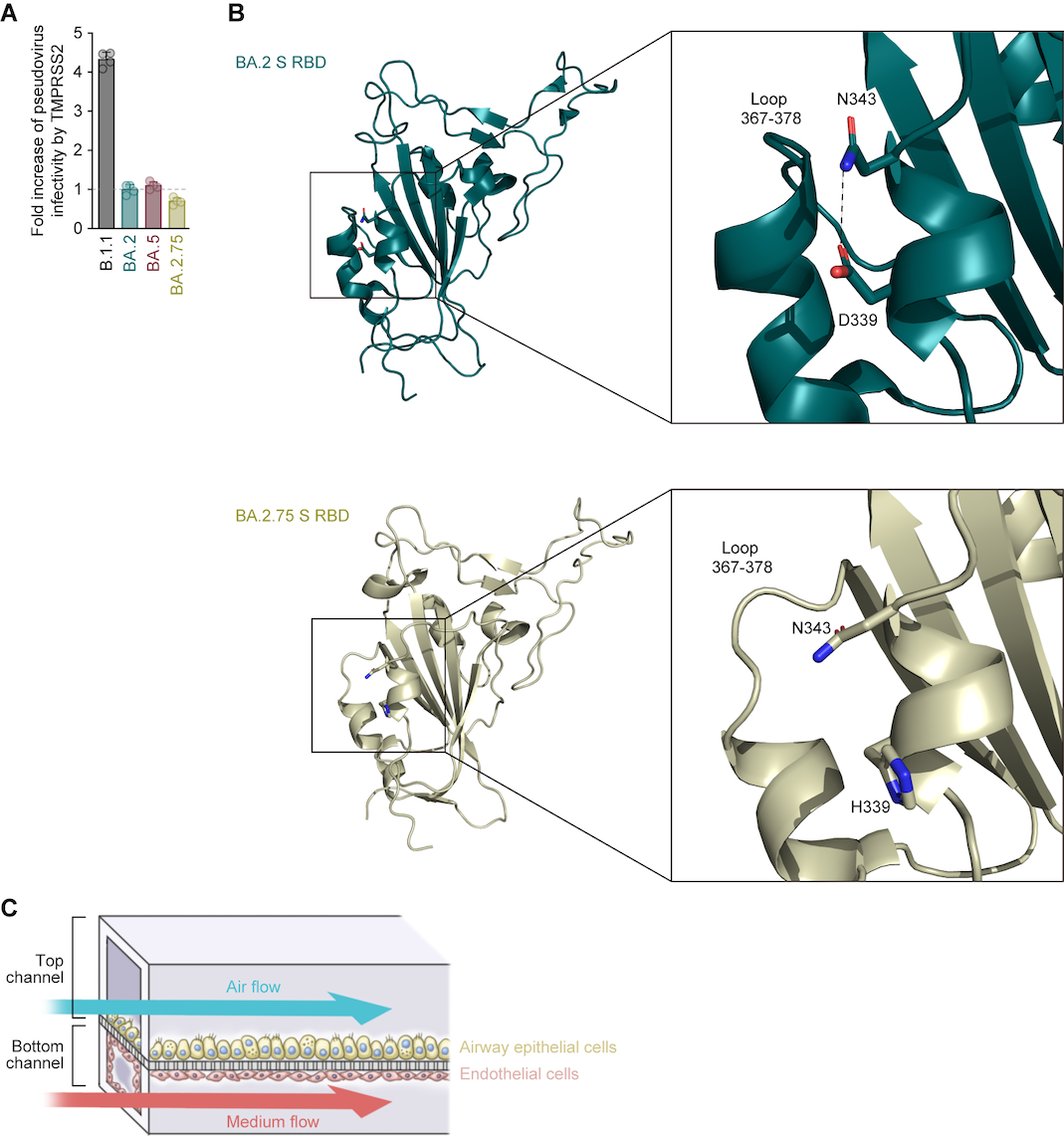

### Figure S4

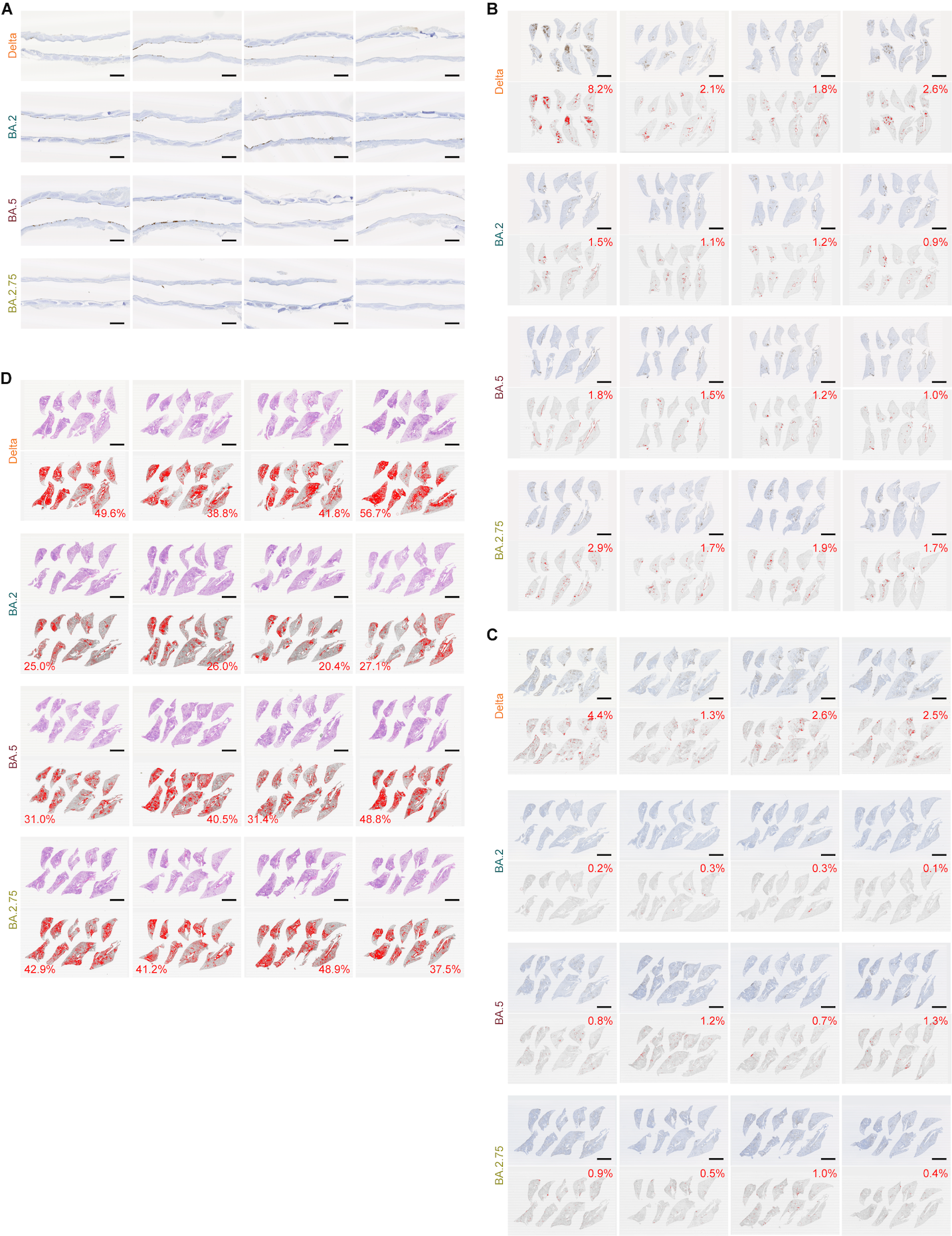
